# Versatile approach for functional analysis of human proteins and efficient stable cell line generation using FLP-mediated recombination system

**DOI:** 10.1101/160101

**Authors:** Roman J. Szczesny, Katarzyna Kowalska, Kamila Klosowska-Kosicka, Aleksander Chlebowski, Ewelina P. Owczarek, Zbigniew Warkocki, Tomasz M. Kulinski, Dorota Adamska, Kamila Affek, Agata Jedroszkowiak, Anna V. Kotrys, Dominik Cysewski, Rafał Tomecki, Pawel S. Krawczyk, Lukasz S. Borowski, Andrzej Dziembowski

## Abstract

Deciphering a function of a given protein requires investigating various biological aspects. Usually, the protein of interest is expressed with a fusion tag that aids or allows subsequent analyses. Additionally, downregulation or inactivation of the studied gene enables functional studies. Development of the CRISPR/Cas9 methodology opened many possibilities but in many cases it is restricted to non-essential genes. It may also be time-consuming if a homozygote is needed. Recombinase-dependent gene integration methods, like the Flp-In system, are very good alternative. The system is widely used in different research areas, which calls for the existence of compatible vectors and efficient protocols that ensure straightforward DNA cloning and creation of stable cell lines. We have created and validated a robust series of 52 vectors for streamlined generation of stable mammalian cell lines using the FLP recombinase-based methodology. Using the sequence-independent DNA cloning method all constructs for a given coding-sequence can be made with just three universal PCR primers. The collection allows tetracycline-inducible expression of proteins with various tags suitable for protein localization, FRET, bimolecular fluorescence complementation (BiFC), protein dynamics studies (FRAP), co-immunoprecipitation, the RNA tethering assay and cell sorting. Some of the vectors contain a bidirectional promoter for concomitant expression of miRNA and mRNA, so that a gene can be silenced and its product replaced by a mutated miRNA-insensitive version. We demonstrate the efficacy of our vectors by creating stable cell lines with various tagged proteins (numatrin, fibrillarin, coilin, centrin, THOC5, PCNA). We have analysed transgene expression over time to provide a guideline for future experiments and compared the utility of commonly used inducers of tetracycline-responsive promoters. We determined the protein interaction network of the exoribonuclease XRN2 and examined the role of the protein in transcription termination by RNAseq analysis of cells devoid of its ribonucleolytic activity. In total we created more than 500 DNA constructs which proves high efficiency of our strategy.

## INTRODUCTION

Deciphering a protein’s function requires investigating its subcellular localisation, identifying its binding partners, and performing multiple functional assays. There are many ways to achieve these goals, with different amounts of required time and effort as well as variable biological relevance of the results obtained. The usual course of action is to express the protein of interest with a fusion tag, a short peptide or a domain, that aids or allows biochemical, cellular or functional analysis. Study of one protein often leads to follow-up experiments that involve other proteins, which can quickly multiply the amount of work required to comprehensively answer the original question. Consequently, straightforward methods or tools that can provide answers to a number of questions are called for.

Ectopic expression is widely used for investigations of human proteins. It can be achieved by transient or stable transfection of cultured cells with a plasmid or virus. Alternatively, one can perform targeted genomic manipulation to engineer the gene of interest in its natural locus. This used to be difficult and time-consuming for most vertebrate cell lines before the advent of CRISPR-based approaches [1, 2, 3]. Genome editing has the crucial advantage in that the studied gene is expressed at its natural levels and naturally responds to all stimuli. However, this approach can prove to be problematic if control of gene expression is required or specific alleles are to be tested. On the other hand, transfection, transient or stable, offers a lot of flexibility in transgene sequence, allowing for the use of stronger, weaker, or even regulated promoters.

Of the two transfection modes – transient and stable – the first one is obviously easier and faster but suffers from cell stress, low reproducibility and heterogeneity of cell populations. Generating stable cell lines eliminates such caveats but requires a considerable amount of time, especially if creating the DNA constructs and cell selection following transfection run into unforeseen problems. These two steps can be streamlined with some careful planning and creating an overarching strategy.

The first step is choosing a reliable cloning method. The traditional one, involving digestion with restriction enzymes and ligation with DNA ligase strongly depends on the target’s sequence; the efficiency of the procedure is highly variable and a universal protocol is virtually impossible. Sequence-independent or recombination based cloning methods like In-Fusion, Gibson assembly, PIPE or SLIC overcome these difficulties [4, 5, 6, 7, 8, 9].

The second step is to use parental cell lines that have been pre-engineered to improve genomic integration of the transgene [10, 11, 12]. One commonly used solution is site-specific recombination employing the FLP recombinase (Figure S1) [13, 14, 15]. In this approach, which is commercially available from Thermo Fisher Scientific as the Flp-In System, the genome of the parental cells contains an FRT sequence (FLP recombination target), which is recognized by the yeast FLP recombinase. The gene of interest is cloned into a plasmid that contains a non-functional antibiotic resistance gene, devoid of a promoter and the initiation codon. Upon FLP-mediated recombination, the plasmid DNA is inserted into the chromosome in such a way that the coding sequence of the new selection marker is substituted for that of the old one. As a result the cells lose one antibiotic resistance and gain another (Figure S1). This guarantees that all selected cells underwent the same integration event and thus all of them have the transgene integrated into the same locus which is not transcriptionally silent. Consequently, clonal selection can be omitted, which greatly simplifies the procedure. Furthermore, monoclonal lines can suffer from occasional genome rearrangements that occur in cells cultured *in vitro*, which could lead do substantial differences between lines [16, 17]; in polyclonal cell lines the effect is diluted across the population.

Several cell lines compatible with the Flp-In system are available [15, 18, 19, 20], including commercial ones. Among them are HeLa and 293 [21, 22, 23, 24], which are the most frequently used non-primary cell lines in basic science and biotechnology [25]. With many reports from wide-ranging topics and detailed transcriptomic and genomic analyses published, HeLa and 293 cells provide a large knowledge base for interpretation of new results [25, 26, 27, 28, 29, 30, 31].

Importantly, expression of the transgene can be driven by inducible promoters, allowing a degree of control over the level, time of expression and its onset. Several inducible systems are used in mammalian cells [32, 33], including ones which respond to ponasteron A (ecdysone analogue) [34, 35, 36], IPTG [37, 38, 39] or tetracycline [40]. The latter utilizes parts of the bacterial tetracycline resistance operon: the repressor protein (TetR) and the operator element found in the promoter of the operon’s structural genes [41]. In a tetracycline-inducible system (Tet-ON) the operator sequence is inserted into the transgene promoter, where it recruits the repressor, resulting in constitutive repression of the transgene, which is alleviated by tetracycline. In an alternative arrangement (Tet-OFF), a mutant TetR is used that can only bind the operator sequence in the presence of tetracycline [42, 43]. Other variants of these systems use fusions of TetR with a transcriptional activator or silencer.

We have established a straightforward method to generate Flp-In-based cellular models for functional studies of human proteins. We have created and validated a robust series of vectors designed for an efficient cloning strategy that enables cheap and easy generation of DNA constructs, and we have combined the efficient cloning with a Flp-In system for stable Tet-ON cell line generation. The cloning procedure has been successfully applied to more than 500 DNA constructs, most of which were obtained in the first attempt. Our vectors facilitate localization studies, protein purification, *in vitro* and *in vivo* protein interaction studies (co-IP, BiFC, FRET), protein dynamics studies (FRAP, photoactivation, etc.) and studies that involve RNA tethering. We present examples of protein localization and interaction studies, including a chemical cross-linking approach that helps to reveal weak interactions. We have analysed transgene expression over time in order to provide a guideline for future experiments. We also compare the utility of two commonly used inducers for tetracycline-responsive promoters, namely tetracycline and doxycycline.

A subset of our vectors enables inducible downregulation of the endogenous gene of interest and concomitant expression of its protein product from an exogenous allele. This approach can be useful *e.g.* for functional validation of newly identified mutations. We used these vectors to establish a cellular model for investigation of the 5’-3’ exoribonuclease XRN2. Deep RNA sequencing analysis of cells devoid of the XRN2 ribonucleolytic activity revealed that this activity is required for transcriptional termination. Our protein interaction studies confirmed previously described XRN2 interactors and revealed potential new ones. Altogether our validation experiments confirm that our strategy is highly applicable for efficient creation of cellular models. Our detailed protocols should ensure smooth transfer of this strategy to other laboratories.

## MATERIALS AND METHODS

### Vector Construction

pKK and pKK-RNAtag vectors were constructed by modifying pcDNA5/FRT/TO (Thermo Fisher Scientific); 5 pKK-BI16, pKK-RNAi, pKK-BiFC and pKK-FRET vectors were constructed by modifying BI16 (a kind gift 1 from Ed Grabczyk), which is in turn derived from pcDNA5/FRT/TO [44]. Standard cloning and SLIC methods were used. Coding sequences of fluorescent proteins and BirA^R118G^ were PCR-amplified from plasmids 5 acquired from Addgene (ID 22010, 22011, 27793, 27795, 27798, 36047, 56172, 62383, 74252, 74279), deposited by: Kyle Roux, Jonathan Weissman, Steven Vogel, Michael Lin, Michael Davidson, Chang-Deng Hu. The MBP tag sequence came from MS2MBP plasmid [45]. The 3xMS2 stem-loop sequence came from 1 pAdML-M3 [46] and the 24xMS2 stem-loop sequence came from pCR24MS2SL – a kind gift of Witold) Filipowicz. The N-terminator peptide sequence came from plasmids described previously [47]. All vector l sequences are presented in Supplementary Data 2 (annotated GenBank format) as well as will be available on our lab’s website, http://adz.ibb.waw.pl (SnapGene format). Vectors will be made available from Addgene.

### Sequence and Ligation Independent Cloning (SLIC)

A simplified SLIC protocol was applied [7]. Detailed description of the SLIC protocol and instructions for 5 using the pKK-RNAi backbone can be found in Supplementary Data 3 and Supplementary Data 4, 7 respectively. Sequences of synthetic DNA (primers and miRNA cassettes) used for construction of plasmids applied in validation experiments are shown in Supplementary Data 5.

### Cell culture

HeLa Flp-In T-REx (a kind gift from Matthias Hentze) and 293 Flp-In T-REx cells (Thermo Fisher Scientific) were cultured in Dulbecco’s Modified Eagle’s Medium (Gibco) supplemented with 10% fetal bovine serum 1 (FBS; standard FBS hereafter) (Gibco) at 37°C in a humidified 5% CO_2_ atmosphere. Where indicated, certified 1 tetracycline-free FBS (Clontech and Biochrom GmbH) was used instead of standard FBS. Cells were tested for mycoplasma contamination.

### Gene Expression Inducers

Tetracyline (550205, Thermo Fisher Scientific) or doxycycline (D9891, Sigma) was added to 96% ethanol at a concentration of 1-5 mg/ml, rotated for 30 minutes at room temperature and incubated overnight at -20°C. On the following day the solutions were rotated again for 30 minutes at room temperature, filtered (0.22 µm) and diluted with ethanol to a final concentration of 0.1 mg/ml. This stock solution (0.1 mg/ml) was stored at -20°C.

### Stable Cell Line Generation

Parental cells were plated onto 6-well plates and cultured for 24 hours. On the next day cells were co-transfected using 2 μl of TransIT-2020 reagent (Mirus) with 0.3 µg of gene-of-interest construct and 1.0 μg of pOG44 (Thermo Fisher Scientific). Twenty four hours after transfection, cells were replated to 60 mm dishes and subjected to selection with hygromycin B (50 and 175 μg/ml for 293 and HeLa cells, respectively) (Thermo Fisher Scientific) and blasticidin (10 μg/ml) (Invivogen) for up to a month. A detailed day-by-day protocol is described in Supplementary Data 6. Where applicable, colonies were stained with crystal violet (0.5% w/v).

### Western Blot

Total protein cell extracts were prepared as described previously [48]. Protein concentration was determined by the Bradford method. The protein extracts, 20 μg per well, were separated by sodium dodecyl sulfate polyacrylamide gel electrophoresis (SDS–PAGE) and transferred to a nitrocellulose membrane (Protran, Whatman GmbH). Western blotting was performed according to standard protocols using the following primary antibodies: anti-EGFP (dilution 1:1000, sc-9666, Santa Cruz Biotechnology), anti-THOC5 (dilution 1:1500, ab86070, Abcam), anti-XRN2 (dilution 1:1000, sc-365258, Santa Cruz Biotechnology). Appropriate horseradish peroxidase-conjugated secondary antibodies (dilution 1:10000, 401393, 401215, Calbiochem) were detected by enhanced chemiluminescence (170-5061, BioRad) according to the manufacturer’s instructions.

### Measurement of Luciferase Activity

293 Flp-In T-REx cells were stably transfected with the BI16 vector. Cells were plated on a 96-wells plate at 5,000 cell per well. Luciferase activity was measured with Dual-Glo Luciferase Assay System (E2920, Promega) according to the manufacturer’s instructions. DTX880 plate reader (Beckman Coulter) was used for measurement of luminescence. Luciferase activity was normalized to the number of cells, which was assessed using AlamarBlue (see next section).

### Resazurin-based Cell Viability Assay

The assay was performed using the AlamarBlue reagent (DAL1100, Thermo Fisher Scientific) according to the manufacturer’s instructions. Briefly, AlamarBlue (1/10 of culture volume) was added to cell culture which was subsequently continued for one hour. Fluorescence was measured with a DTX880 plate reader (Beckman Coulter) using 535/25 and 595/35 filters (excitation and emission, respectively).

### Flow Cytometry

#### Expression kinetics

Cells were plated onto 24-well plates at 50,000 cells per well. 24 hours later the cells were induced with tetracycline or doxycycline at a final concentration of 100 ng/ml in 6 hour intervals. Additionally, at the time of the first induction another well was plated, where the cells were induced upon plating. 24 hours after the first induction the cells were detached by trypsin treatment, diluted with PBS, and analysed with a BD FACSCalibur flow cytometer (BD Biosciences). A gate was applied to the FSC/SSC plot to exclude dead cells and debris. 10,000 events were collected. Data were analysed using the FlowingSoftware.

#### Validation of XRN2 stable cell lines

Cells were plated onto 6-well plates at 500,000 cells per well and induced upon plating with tetracycline (25 ng/ml). 24 hours later the cells were detached by trypsin treatment, washed with PBS, and analysed with an Attune NxT flow cytometer equipped with 488 nm and 561 nm laser diodes (Thermo Fisher Scientific). A gate was applied to the FSC/SSC plot to exclude dead cells and debris. Doublet discrimination was performed based on the FSC-A/FSC-H plot. 20,000 events were collected. Data were analysed using Attune NxT software.

### Fluorescence Microscopy

Stable cell lines expressing XRN2 were analysed by the following protocol: 24 hours prior to imaging cells were seeded on poly-L-lysine-coated (see below) 8-well Lab-Tek II Chambered Coverglass culture vessels (155409, Thermo Fisher Scientific) at 30,000 per well and induced with tetracycline (25 ng/ml). Before imaging, Hoechst 33342 dye was added to the medium (2 ng/ml) for 30 minutes to stain cell nuclei. After staining medium was replaced. Images were collected using a FluoView1000 Olympus confocal system with a PLANAPO 60x/1.40 oil immersion lens. Live cell imaging was performed in a temperature (37°C) and CO_2_ (5%) incubator.

All other stable cell lines were analyzed as follows: cells were plated onto 8-well Lab-Tek II Chambered Coverglass culture vessels coated with poly-L-lysine at 7,000 (HeLa) or 10,000 (293) cells per well. Tetracycline at a final concentration of 25 ng/ml was added to the medium upon plating. On the following day the cells were stained with Hoechst 33342 for 30 minutes (final concentration of 50 ng/ml). Live imaging was done with an FV10i system (Olympus), maintaining the cells at 37°C in a humidified 5% CO_2_ atmosphere delivered from a The Brick gas mixer (Life Imaging Services). Fluorescence was excited with 405 and 483 laser diodes and collected with a SuperApochromat 60x/1.2 water immersion lens. Where applicable, Z stacks were collected with 350 nm spacing.

### Preparation of poly-L-lysine-coated coverglasses

Poly-L-lysine hydrobromide (P1274, Sigma) was dissolved in sterile water to 0.01% (w/v). The solution was sterilized with a 0.22 micron filter before freezing aliquots at -20°C. Before use, the solution was thawed and 200 μl was added to each well of 8-well Lab-Tek II Chambered Coverglass to fully coat the surface of each well. Coating was performed for 1 hour at 37°C, after that the solution was removed and coverglasses were dried at room temperature for 20 minutes under a laminar flow hood. The poly-L-lysine solution was collected and stored at -20°C for repeated use. Dried coverglasses were stored at room temperature or directly used for cell seeding.

### RNA Isolation, Library Construction and Deep-Sequencing

Total RNA was isolated with the TRI Reagent (T9424, Sigma) according to the manufacturer’s instructions. DNA contamination from 2 μg of nucleic acids was removed by 2 U of TURBO DNase (AM2238, Ambion) in 20 µl of the supplied buffer in 37°C for 30 min. RNA was extracted with phenol-chloroform, precipitated with ethanol and resuspended in RNase free water. Concentration was measured with NanoDrop 2000 Spectrophotometer (Thermo Fisher Scientific). Prior to library preparation, to provide an internal performance control for further steps, 1 μg of RNA was mixed with 4 μl of 1:99 diluted ERCC RNA Spike-In Control Mix 1 (4456740, Ambion). Subsequently, rRNA was depleted using Ribo-Zero Gold rRNA Removal Kit (MRZG12324, Human/Mouse/Rat, Illumina) according to the manufacturer’s protocol.

RNA-seq libraries were constructed as previously described in Sultan et al., 2012 [49] with minor modifications. Fragmentation and first strand cDNA synthesis were performed as in TruSeq RNA Library Prep kit v2 protocol (Illumina, RS-122-2001, instruction number 15026495 Rev. D), using SuperScript III reverse transcriptase (18080-085, Thermo Fisher Scientific). For second strand synthesis, reaction mixtures were supplemented with 1 μl of 5x First Strand Synthesis Buffer (18080-085, Thermo Fisher Scientific), 15 μl 5x Second Strand Synthesis Buffer (10812-014, Thermo Fisher Scientific), 0.45 μl 50 mM MgCl, 1 μl 100 mM DTT, 2 μl of 10 mM dUNTP Mix (dATP, dGTP, dCTP, dUTP, 10 mM each, R0182, R0133, Thermo Fisher Scientific), water to 57 μl, 5 U *E. coli* DNA Ligase (M0205L, NEB), 20 U *E. coli* DNA Polymerase I (NEB, M0209L), 1 U RNase H (18021-071, Thermo Fisher Scientific), and incubated at 16°C for 2h. Further steps: purification, end-repair, A-tailing and adapter ligation were performed as described in the previously mentioned TruSeq kit protocol with one modification: the first purification eluate was not decanted from the magnetic beads and subsequent steps were performed with the beads in solution. Instead of a new portion of magnetic beads, an equal volume of 20% PEG 8000 in 2.5 M NaCl was added and the DNA bound to the beads already present in the mixture. After the second clean up procedure after adapter ligation, the supernatant was separated from the beads and treated with USER Enzyme (M5505L, NEB) in 1x UDG Reaction Buffer (M0280S, NEB) at 37°C for 30 min. The digestion step ensures that the second strand synthetized with dUTP instead of dTTP is removed from cDNA, resulting in strand-specific libraries. The product was amplified using 1 U of Phusion High-Fidelity DNA Polymerase (F530L, Thermo Fisher Scientific) in 1x HF Buffer supplemented with 0.2 mM dNTP Mix, and the following primers: PP1 (5’-AATGATACGGCGACCACCGAGATCTACACTCTTTCCCTACACGA-3’), PP2 (5’-CAAGCAGAAGACGGCATACGAGAT-3’). TruSeq kit protocol temperature scheme with 12 amplification cycles and subsequent purification procedure was applied. Enriched library quality was verified using 2100 Bioanalyzer and High Sensitivity DNA kit (5067-4626, Agilent). The libraries’ concentration was estimated by qPCR means with KAPA Universal Library Quantification Kit (KK4824, Kapa Biosystems), according to the supplied protocol. Sequencing was carried out on an Illumina NextSeq 500 sequencing platform, using NextSeq 500 High Output Kit (150 cycles) (FC-404-1002, Illumina) and standard libraries denaturation and pair-end sequencing procedures (Instructions: 15048776 Rev. D, 15046563 Rev. F) of 2×75 cycles.

### Analysis of Deep Sequencing Data

Strand-specific RNAseq libraries (dUTP RNA) were prepared in triplicate for each condition and sequenced in the 75-nt paired-end mode to the average depth of 10 million reads (GEO accession number: GSE99421, security token: ghmheeqytvkfbof). Reads were mapped to the reference human genome (GRCh38) using the STAR short read aligner [50]. Quality control, read processing and filtering, visualization of the results and counting of reads for the Genecode v22 comprehensive annotation were performed using custom scripts using elements of the RSeQC, BWtools, BEDtools and SAMtools packages. Transcripts were annotated using StringTie [51]. The merged unmodified 293 Flp-In T-REx cells annotation was used to perform meta-gene analysis of the transcriptional read-through in wild-type and mutant XRN2 cells. Cumulative, strand-specific signal was centered around 3’ ends of highly expressed (TMP > 10), spliced transcripts and normalized to the signal within the last 250 nt of the analyzed transcripts.

### DSP Cross-linking

20 mM DSP (22586, Thermo Fisher Scientific) stock solution in DMSO was prepared directly before use. 293 Flp-In T-REx cells (1×10^6^ per well of a 6-well plate) were plated 24 hours before treatment. Cells were washed with PBS and incubated in PBS containing different concentrations of DSP ranging from 0.05 mM to 1.0 mM for 15 or 45 minutes at room temperature with gentle mixing. The cross-linking reaction was terminated by adding Tris-HCl pH 7.5 to 50 mM and incubating at room temperature for 15 minutes with gentle mixing. Cells were then detached by pipetting, centrifuged at 400g for 5 minutes at room temperature, resuspended in PBS and centrifuged as previously. Supernatant was discarded and pelleted cells were lysed in RIPA buffer (10 mM Tris-HCl pH 7.4, 140 mM NaCl, 5 mM EDTA, 1% (v/v) Triton X-100, 1% (w/v) deoxycholate, 0,1% (w/v) SDS) containing a protease inhibitor cocktail. 20 μg of lysates were separated in an SDS-PAGE gel and transferred to a nitrocellulose membrane. Loading buffer without 2-mercaptoethanol was applied, if not otherwise stated. A standard protocol for western blotting was applied (see above). Anti-HSP60 antibodies (Santa Cruz Biotechnology: sc-1722) were used.

### Protein Purification

293 Flp-In T-REx cells or their derivatives expressing tagged XRN2 for 24 hours were treated with DSP or not, washed with PBS, collected and stored at -80°C. Material from six 145 mm plates was used for each purification condition. Cells were thawed in lysis buffer (150 mM NaCl, 10 mM Tris-HCl pH 8.0, 1% (v/v) Triton X-100, benzonase (50 U/ml, E1014, Sigma), protease inhibitor cocktail), incubated for 10 minutes on ice and subjected to sonication (20 cycles, 30 s ON / 30 s OFF, high mode, Bioruptor XL, Diagenode). Lysates were centrifuged for 15 minutes at 12,000g at 4°C. Protein extracts were mixed with beads coupled with recombinant anti-GFP antibodies [52] (40 μl of beads volume was used per purification) and rotated for 1.5 h at 4°C. Beads were washed with 20 volumes of washing buffer A (20 mM HEPES-KOH 7.1, 10% (v/v) glycerol, 3 mM MgCl_2_, 0.5% (v/v) Igepal CA630, protease inhibitor cocktail, 150 mM or 300 mM NaCl for DSP untreated or treated samples, respectively) and 10 volumes of washing buffer B (150 mM NaCl, 10 mM Tris-HCl pH 8.0). In the case of DSP-treated samples purified proteins were eluted by incubation of beads in 3.5 volumes of pre-heated elution buffer (3% (w/v) SDS, 50 mM Tris-HCl pH 8.0, 50 mM DTT) for 10 minutes at 90°C. For DSP-untreated samples purified proteins were eluted with TEV protease. Beads were resuspended in 3.5 volumes of TEV cleavage buffer (150 mM NaCl, 0.5 mM EDTA, 10 mM Tris-HCl pH 8.0, 1 mM DTT) containing TEV protease purified in our laboratory [52] and incubated for 2 hours at room temperature. Regardless of elution method, eluted proteins were precipitated as described in the next section.

### Protein Precipitation

Proteins were precipitated using the methanol-chloroform method [53]. The elution fraction was mixed with 4 volumes of methanol, briefly vortexed, and mixed with one volume of chloroform, after which samples were briefly vortexed and 3 volumes of water were added. After vortexing samples were centrifuged for 2 minutes at 14,000g at 22°C. The aqueous phase was discarded and 4 volumes of methanol were added. Samples were i briefly vortexed and centrifuged for 3 minutes at 14,000g at 22°C. The supernatant was discarded and the pellet was dried at room temperature and stored at -20°C.

### Mass spectrometry analysis

Precipitated proteins were dissolved in 100 μl of 100 mM ammonium bicarbonate buffer, reduced in 100 mM DTT for 30 min at 57°C, alkylated in 55 mM iodoacetamide for 40 min at RT in the dark and digested overnight with 10 ng/ml trypsin (V5280, Promega) at 37°C. Finally, to stop the digestion, trifluoroacetic acid was added at a final concentration of 0.1%. The mixture was centrifuged at 4°C, 14,000g for 20 min to remove precipitated material. MS analysis was performed by LC-MS in the Laboratory of Mass Spectrometry (IBB PAS, Warsaw) using a nanoAcquity UPLC system (Waters) coupled to a LTQ-Orbitrap Velos or QExactive’ mass spectrometer (Thermo Fisher Scientific). The mass spectrometer was operated in the data-dependent MS2 mode, and data were acquired in the m/z range of 300-2000. Peptides were separated by a 180 min linear gradient of 95% solution A (0.1% formic acid in water) to 35% solution B (0.1% formic acid in acetonitrile). The measurement of each sample was preceded by three washing runs to avoid cross-contamination. The final MS washing run was searched for the presence of cross-contamination between samples. If the protein of interest was identified in the washing run and in the next measured sample at the same or smaller intensity, then the identified protein was treated as cross-contamination. Data were analyzed with the Max-Quant (Version 1.4.1.2) platform using mode match between runs [54]. The reference human proteome database from UniProt was used (downloaded at 2015.11.20). Variable modification were set for: methionine oxidation, carbamidomethyl on cysteines and reduced or alkylated part of DSP spacer arm in the case of DSP-treated sample. Protein abundance was defined as the signal intensity calculated by MaxQuant software for a protein divided by its molecular weight. Enrichment was defined as the ratio of the protein intensity measured in the bait purification to background level (*i.e.*, protein intensity in the negative control purification with the background level arbitrarily set to 1 for proteins not detected in the negative control).

## RESULTS AND DISCUSSION

### Construction of SLIC-competent vectors

A new set of vectors was designed to fulfil four requirements: 1) compatibility with sequence-independent, straightforward and efficient cloning; 2) minimal number of primers required for cloning into different vectors; 3) compatibility with FLP-mediated stable cell line generation; 4) regulated transgene expression. To achieve these goals we modified the pcDNA5/FRT/TO vector (Thermo Fisher Scientific) that enables FLP-mediated cell line generation and contains a tetracycline-regulated CMV promoter to drive expression of the cloned CDS (Figure S1). The aim of our modification was to make the vector suitable for the SLIC approach (sequence and ligation independent cloning) [4]. The multiple cloning site of the vector was modified by introducing two 21-nucleotide long sequences, the SLIC arms (Figure 1). These fragments are complementary to the ends of the linear DNA fragment to be cloned. They encompass TEV-L and TEV-R sequences which encode a 7 amino acid long peptide which is recognized by the tobacco etch virus (TEV) protease [55] (Figure 1A).

**Figure 1.**
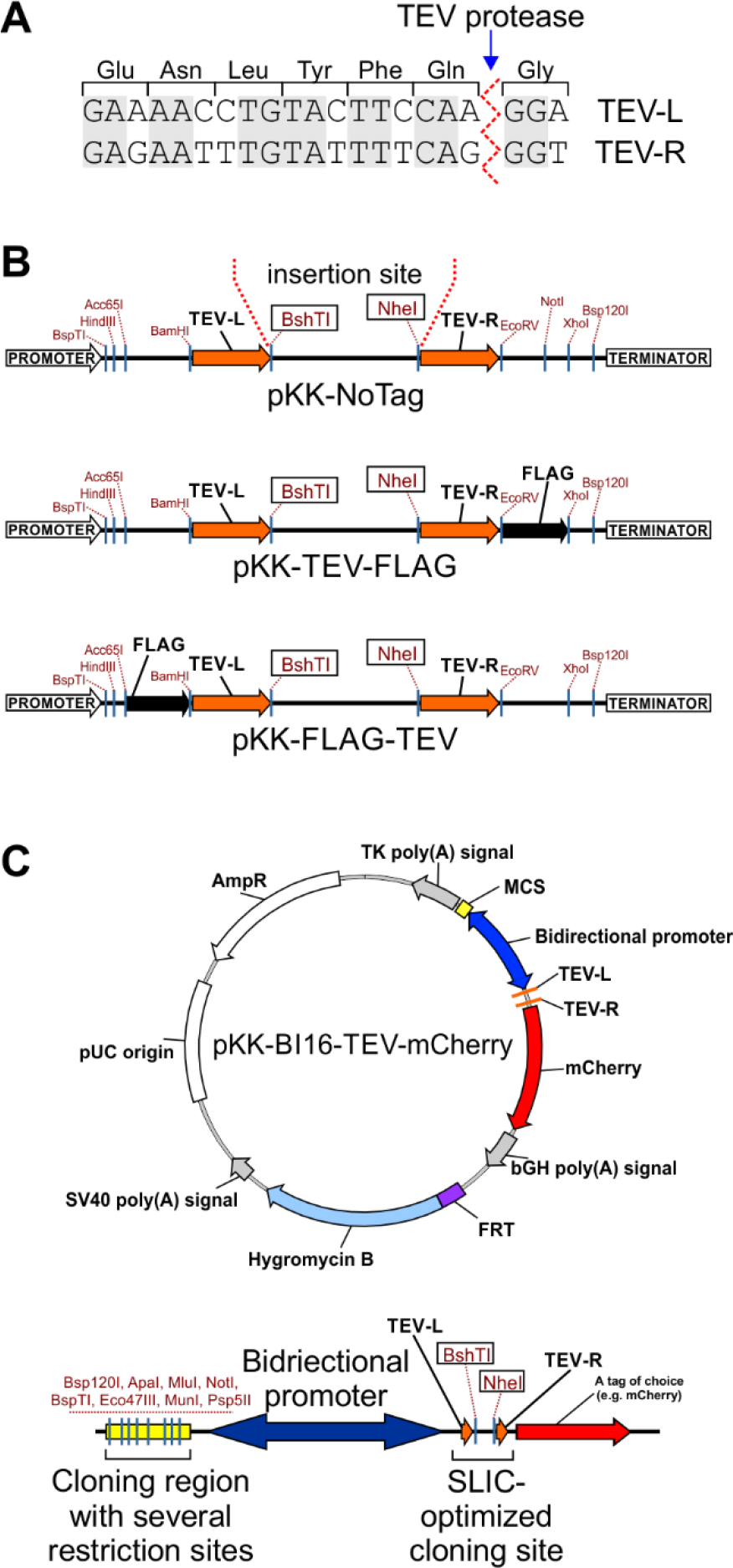
The pKK vector series. (A) Nucleotide sequences of the TEV-L and TEV-R. Translation to protein and TEV protease cleavage site are shown. Shaded letters indicate nucleotides common to both sequences. (B) Cloning sites of selected pKK vectors. Potentially useful unique restriction sites are marked. For all pKK vectors BshTI and NheI restriction enzymes are used for vector linearization before DNA cloning with the help of our universal SLIC protocol. (C) Example of a pKK-BI16 vector. Map of pKK-BI16-TEV-mCherry vector and its cloning region (bottom diagram). Useful unique restriction sites are marked.

Depending on the vector, the SLIC arms lie upstream or downstream of a tag coding sequence (Figure 1B; maps of all reported vectors are presented in Supplementary Data 7). Several vectors with different tags were constructed: 1) short tags, *e.g.* FLAG; 2) fluorescent proteins, *e.g.* EGFP; 3) humanized biotin ligase – BirA, which can be used for identification of proximal and interacting proteins; 4) other proteins, *e.g.* protein A or maltose binding protein (MBP), for which several molecular tools are described (vectors pKK, Table 1 and Supplementary Data 8). The coding sequence of the protein of interest is inserted between the SLIC arms. For the N-terminal fusions the initiation codon originates from the vector and the termination codon is introduced within the cloned fragment, and vice-versa in the case of C-terminal fusions. The same forward primer can be used to create both N- and C-terminal fusions, however, a specific reverse primer is required (with or without the stop codon). Thus, only 3 primers are sufficient to prepare constructs encoding a protein of interest in N- or C-terminal fusion with different tags and each tag encoded by pKK-series vectors can be cleaved off with the TEV protease. Importantly, synonymous codons were used for designing of TEV-L and TEV-R sequences to decrease chance of intramolecular homologous DNA recombination.

**Table 1.**
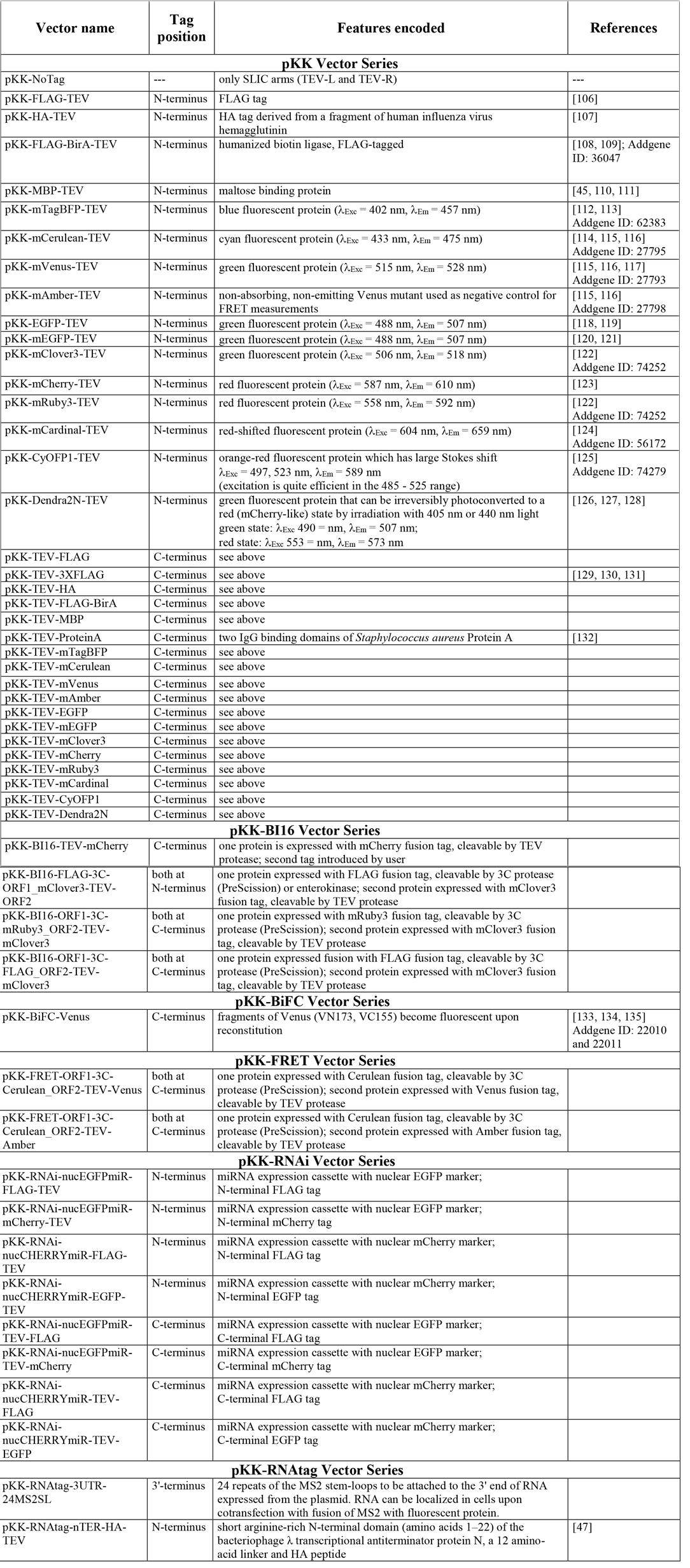
List of created vectors. Full description of the vectors can be found in and Supplementary Data 8. For all vectors BshTI and NheI restriction enzymes are used for vector linearization before SLIC cloning according to our universal protocol.

| Vector name | Tag position | Features encoded | References |
| --- | --- | --- | --- |
| pKK Vector Series |  |  |  |
| pKK-NoTag | --- | only SLIC arms (TEV-L and TEV-R) | --- |
| pKK-FLAG-TEV | N-terminus | FLAG tag | [106] |
| pKK-HA-TEV | N-terminus | HA tag derived from a fragment of human influenza virus hemagglutinin | [107] |
| pKK-FLAG-BirA-TEV | N-terminus | humanized biotin ligase, FLAG-tagged | [108, 109]; Addgene ID: 36047 |
| pKK-MBP-TEV | N-terminus | maltose binding protein | [45, 110, 111] |
| pKK-mTagBFP-TEV | N-terminus | blue fluorescent protein ($\lambda_{Exc} = 402$ nm, $\lambda_{Em} = 457$ nm) | [112, 113]<br>Addgene ID: 62383 |
| pKK-mCerulean-TEV | N-terminus | cyan fluorescent protein ($\lambda_{Exc} = 433$ nm, $\lambda_{Em} = 475$ nm) | [114, 115, 116]<br>Addgene ID: 27795 |
| pKK-mVenus-TEV | N-terminus | green fluorescent protein ($\lambda_{Exc} = 515$ nm, $\lambda_{Em} = 528$ nm) | [115, 116, 117]<br>Addgene ID: 27793 |
| pKK-mAmber-TEV | N-terminus | non-absorbing, non-emitting Venus mutant used as negative control for FRET measurements | [115, 116]<br>Addgene ID: 27798 |
| pKK-EGFP-TEV | N-terminus | green fluorescent protein ($\lambda_{Exc} = 488$ nm, $\lambda_{Em} = 507$ nm) | [118, 119] |
| pKK-mEGFP-TEV | N-terminus | green fluorescent protein ($\lambda_{Exc} = 488$ nm, $\lambda_{Em} = 507$ nm) | [120, 121] |
| pKK-mClover3-TEV | N-terminus | green fluorescent protein ($\lambda_{Exc} = 506$ nm, $\lambda_{Em} = 518$ nm) | [122]<br>Addgene ID: 74252 |
| pKK-mCherry-TEV | N-terminus | red fluorescent protein ($\lambda_{Exc} = 587$ nm, $\lambda_{Em} = 610$ nm) | [123] |
| pKK-mRuby3-TEV | N-terminus | red fluorescent protein ($\lambda_{Exc} = 558$ nm, $\lambda_{Em} = 592$ nm) | [122]<br>Addgene ID: 74252 |
| pKK-mCardinal-TEV | N-terminus | red-shifted fluorescent protein ($\lambda_{Exc} = 604$ nm, $\lambda_{Em} = 659$ nm) | [124]<br>Addgene ID: 56172 |
| pKK-CyOFP1-TEV | N-terminus | orange-red fluorescent protein which has large Stokes shift<br>$\lambda_{Exc} = 497, 523$ nm, $\lambda_{Em} = 589$ nm<br>(excitation is quite efficient in the 485 - 525 range) | [125]<br>Addgene ID: 74279 |
| pKK-Dendra2N-TEV | N-terminus | green fluorescent protein that can be irreversibly photoconverted to a red (mCherry-like) state by irradiation with 405 nm or 440 nm light<br>green state: $\lambda_{Exc} 490 =$ nm, $\lambda_{Em} = 507$ nm;<br>red state: $\lambda_{Exc} 553 =$ nm, $\lambda_{Em} = 573$ nm | [126, 127, 128] |
| pKK-TEV-FLAG | C-terminus | see above |  |
| pKK-TEV-3XFLAG | C-terminus | see above | [129, 130, 131] |
| pKK-TEV-HA | C-terminus | see above |  |
| pKK-TEV-FLAG-BirA | C-terminus | see above |  |
| pKK-TEV-MBP | C-terminus | see above |  |
| pKK-TEV-ProteinA | C-terminus | two IgG binding domains of *Staphylococcus aureus* Protein A | [132] |
| pKK-TEV-mTagBFP | C-terminus | see above |  |
| pKK-TEV-mCerulean | C-terminus | see above |  |
| pKK-TEV-mVenus | C-terminus | see above |  |
| pKK-TEV-mAmber | C-terminus | see above |  |
| pKK-TEV-EGFP | C-terminus | see above |  |
| pKK-TEV-mEGFP | C-terminus | see above |  |
| pKK-TEV-mClover3 | C-terminus | see above |  |
| pKK-TEV-mCherry | C-terminus | see above |  |
| pKK-TEV-mRuby3 | C-terminus | see above |  |
| pKK-TEV-mCardinal | C-terminus | see above |  |
| pKK-TEV-CyOFP1 | C-terminus | see above |  |
| pKK-TEV-Dendra2N | C-terminus | see above |  |
| pKK-BI16 Vector Series |  |  |  |
| pKK-BI16-TEV-mCherry | C-terminus | one protein is expressed with mCherry fusion tag, cleavable by TEV protease; second tag introduced by user |  |
| pKK-BI16-FLAG-3C-ORF1_mClover3-TEV-ORF2 | both at N-terminus | one protein expressed with FLAG fusion tag, cleavable by 3C protease (PreScission) or enterokinase; second protein expressed with mClover3 fusion tag, cleavable by TEV protease |  |
| pKK-BI16-ORF1-3C-mRuby3_ORF2-TEV-mClover3 | both at C-terminus | one protein expressed with mRuby3 fusion tag, cleavable by 3C protease (PreScission); second protein expressed with mClover3 fusion tag, cleavable by TEV protease |  |
| pKK-BI16-ORF1-3C-FLAG_ORF2-TEV-mClover3 | both at C-terminus | one protein expressed fusion with FLAG fusion tag, cleavable by 3C protease (PreScission); second protein expressed with mClover3 fusion tag, cleavable by TEV protease |  |
| **pKK-BiFC Vector Series** |  |  |  |
| pKK-BiFC-Venus | C-terminus | fragments of Venus (VN173, VC155) become fluorescent upon reconstitution | [133, 134, 135]<br>Addgene ID: 22010 and 22011 |
| **pKK-FRET Vector Series** |  |  |  |
| pKK-FRET-ORF1-3C-Cerulean_ORF2-TEV-Venus | both at C-terminus | one protein expressed with Cerulean fusion tag, cleavable by 3C protease (PreScission); second protein expressed with Venus fusion tag, cleavable by TEV protease |  |
| pKK-FRET-ORF1-3C-Cerulean_ORF2-TEV-Amber | both at C-terminus | one protein expressed with Cerulean fusion tag, cleavable by 3C protease (PreScission); second protein expressed with Amber fusion tag, cleavable by TEV protease |  |
| **pKK-RNAi Vector Series** |  |  |  |
| pKK-RNAi-nucEGFPmiR-FLAG-TEV | N-terminus | miRNA expression cassette with nuclear EGFP marker; N-terminal FLAG tag |  |
| pKK-RNAi-nucEGFPmiR-mCherry-TEV | N-terminus | miRNA expression cassette with nuclear EGFP marker; N-terminal mCherry tag |  |
| pKK-RNAi-nucCHERRYmiR-FLAG-TEV | N-terminus | miRNA expression cassette with nuclear mCherry marker; N-terminal FLAG tag |  |
| pKK-RNAi-nucCHERRYmiR-EGFP-TEV | N-terminus | miRNA expression cassette with nuclear mCherry marker; N-terminal EGFP tag |  |
| pKK-RNAi-nucEGFPmiR-TEV-FLAG | C-terminus | miRNA expression cassette with nuclear EGFP marker; C-terminal FLAG tag |  |
| pKK-RNAi-nucEGFPmiR-TEV-mCherry | C-terminus | miRNA expression cassette with nuclear EGFP marker; C-terminal mCherry tag |  |
| pKK-RNAi-nucCHERRYmiR-TEV-FLAG | C-terminus | miRNA expression cassette with nuclear mCherry marker; C-terminal FLAG tag |  |
| pKK-RNAi-nucCHERRYmiR-TEV-EGFP | C-terminus | miRNA expression cassette with nuclear mCherry marker; C-terminal EGFP tag |  |
| **pKK-RNAtag Vector Series** |  |  |  |
| pKK-RNAtag-3UTR-24MS2SL | 3'-terminus | 24 repeats of the MS2 stem-loops to be attached to the 3' end of RNA expressed from the plasmid. RNA can be localized in cells upon cotransfection with fusion of MS2 with fluorescent protein. |  |
| pKK-RNAtag-nTER-HA-TEV | N-terminus | short arginine-rich N-terminal domain (amino acids 1–22) of the bacteriophage $\lambda$ transcriptional antiterminator protein N, a 12 amino-acid linker and HA peptide | [47] |

To extend the variety of possible functional analyses we also devised a vector with a bidirectional promoter which enables simultaneous, inducible expression of two introduced genes (Figure 1C). To this end we used the BI16 plasmid [44], a derivative of pcDNA5/FRT/TO with the original CMV promoter duplicated back-to-back. The tetracycline operator sequence was maintained so that transcription in both directions is regulated by the tetracycline repressor. We removed the luciferase coding sequences from BI16 and redesigned the cloning sites. As a result, we created a series of pKK-BI16 vectors (Table 1 and Supplementary Data 8). They utilize the cloning strategy described above on one side of the bidirectional promoter and have a traditional multiple cloning site on the other (Figure 1C, bottom diagram). These vectors should significantly simplify construction of plasmids for expression of two independent coding sequences. The pKK-BI16 vectors were further modified to obtain the pKK-RNAi, pKK-BiFC, and pKK-FRET vector series (Table 1 and Supplementary Data 8), which are intended for functional studies and *in vivo* protein interaction studies using bimolecular fluorescence complementation or Förster resonance energy transfer approach.

### Efficiency of the cloning procedure

The cloning procedure starts with PCR amplification of a DNA fragment to be cloned using primers with overhangs complementary to the SLIC arms in the vector (Figure 2). The PCR product has to be purified from nucleotides, which inhibit the subsequent SLIC reaction. The purification method depends on the specificity of the PCR reaction; all unspecific PCR products must be removed. After purification the product is mixed with a linearized vector and treated with T4 DNA polymerase. In the absence of nucleotides the polymerase trims 3’ ends of DNA, producing sticky ends both in the vector and the PCR product (Figure 2). The sticky ends hybridize and form a nicked, potentially gapped, DNA molecule. After introduction to bacteria such lesions are repaired by the host system (Figure 2).

**Figure 2.**
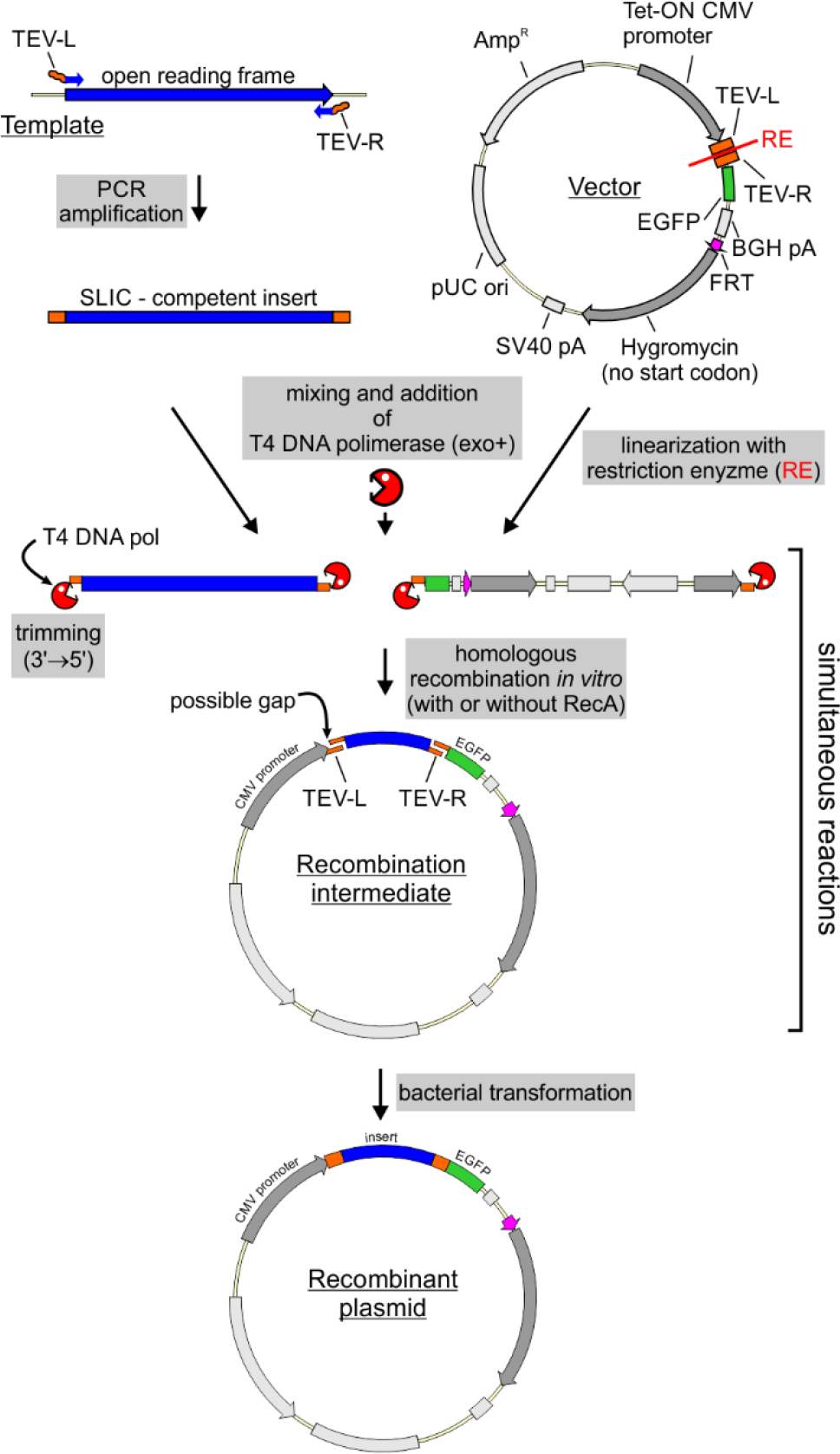
SLIC-based DNA cloning strategy. See main text for detailed description. RE - restriction enzymes used for vector linearization. These are BshTI and NheI in our protocol for universal SLIC. EGFP is an example of tag which can be used.

In the course of our studies we used this strategy to clone 155 different protein coding sequences to pKK vectors series, some fused with different tags, bringing the total number of constructed plasmids to 456, a number high enough to assess the efficiency of our strategy (Table 2). In the analysis of cloning efficiency we considered the PCR template (plasmid versus cDNA) and the number of attempts required to get designed construct. The first attempt was successful in 99% of cases with a plasmid template and in 75% of cases with a cDNA template (Table 2). We were unable to obtain 16% of designed constructs. This failure resulted mostly from unsuccessful PCR amplification of the insert (59 cases out of 72 unsuccessful clonings). Thus, some constructs may require additional optimization steps to produce the insert. Nevertheless, the analysis demonstrates the high efficiency of our strategy and points to insert preparation as the limiting step of the procedure.

**Table 2.** SLIC cloning efficiency. Efficiency of CDS cloning into pKK vector series using our universal SLIC protocol.

|  |  | Constructs designed | Successful first attempt | Unsuccessful cloning due to problems in insert PCR amplification | Unsuccessful cloning in total | First attempt cloning rate |
| --- | --- | --- | --- | --- | --- | --- |
| Template | cDNA | 281 | 211 | 59 | 70 | 75% |
|  | Plasmid | 175 | 173 | 2 | 2 | 99% |
|  | Total | 456 | 384 | 61 | 72 | 84% |

### New vectors are competent for stable cell line generation

We set out to examine whether the modifications to the pcDNA5/FRT/TO backbone interfere with stable cell line generation. In order to establish a baseline for FLP-mediated integration efficiency, we transfected the 293 Flp-In T-REx parental cell line with the original pcDNA5/FRT/TO vector and counted the number of colonies that originated from cells that had undergone FRT-targeted plasmid integration and became resistant to the selection antibiotic. We tested a range of selection antibiotics concentrations as well as the amounts of the targeting vector used for transfection (Figure 3A). We found that decreasing the antibiotic concentration yields more colonies, while providing enough selective pressure to kill off non transfected cells (Figure 3A). As for DNA amount, we found that within the examined range it had no effect on the number of colonies obtained (Figure 3A).

Having optimized transfection and selection conditions, we compared stable transfection efficiency between the original vector and our vectors bearing N- or C-terminal EGFP tags (pKK-EGFP-TEV and pKK-TEV-EGFP, respectively). We found that our vectors produced a lower number of colonies (Figure 3B), however, this number is enough to establish functional stable cell lines, as evidenced by our further experiments (see below). We also observed that the number of colonies obtained for a given plasmid can vary significantly from transfection to transfection (72 versus 46 colonies for pcDNA5/FRT/TO in Figure 3A and 3B, respectively), which was not due to any obvious reasons like different plasmid preparations or number of cells subjected to transfection.

**Figure 3.**
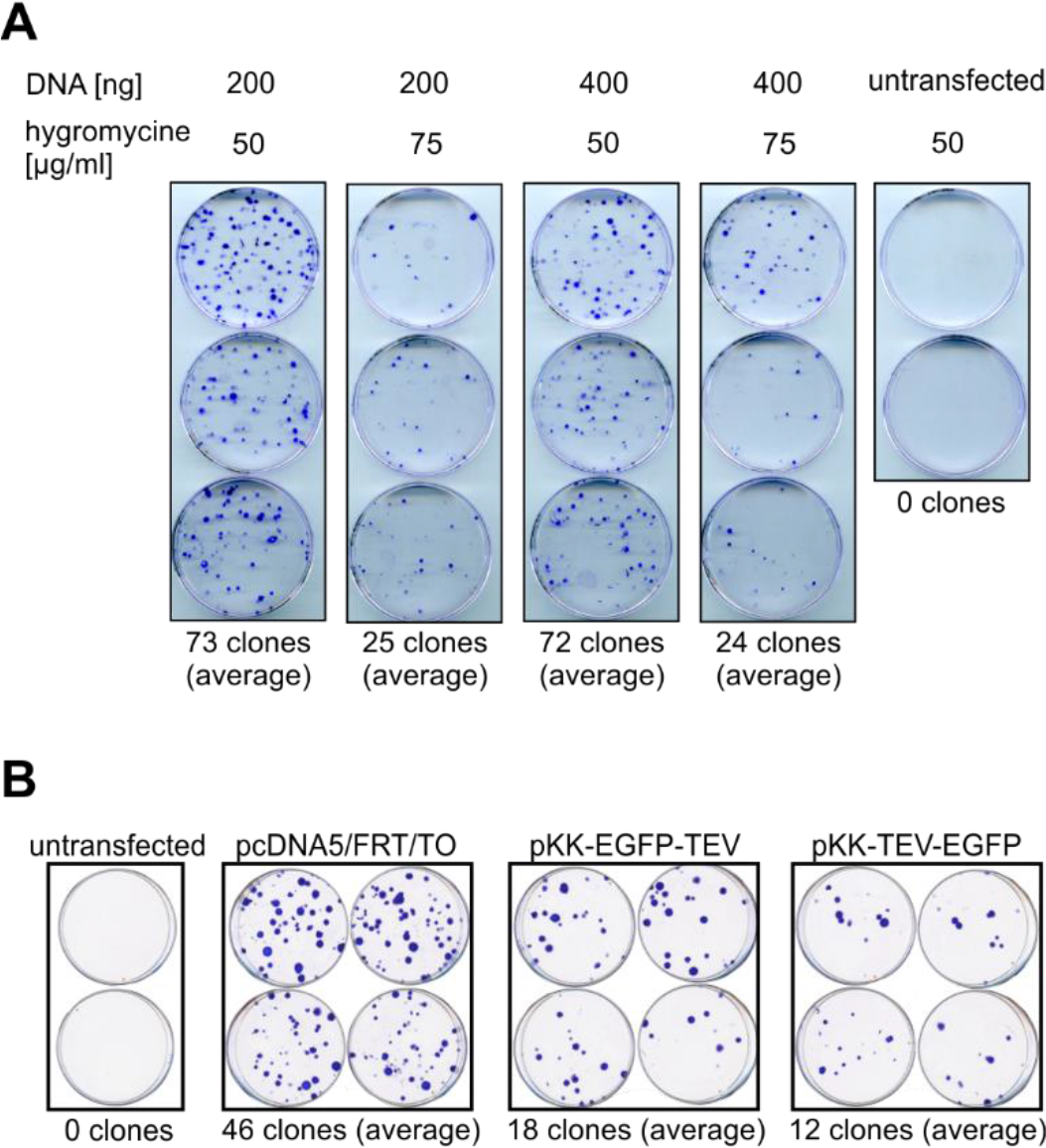
Efficiency of stable cell line generation. (A) Influence of plasmid quantity and selection stringency on the number of colonies obtained following stable transfection of 293 Flp-In T-REx cells. 1.0 μg of pOG44 was mixed with the indicated amounts of pcDNA5/FRT/TO and used for transfection. Cells were selected by treatment with the indicated concentration of hygromycin B and constant concentration of blasticidin S (10 pg/ml). Colonies were stained with crystal violet. (B) Comparison of stable transfection efficiency with pcDNA5/FRT/TO or its pKK derivatives. Cells were transfected with 300 ng of indicated plasmids and 1.0 μg of pOG44 and subjected to selection with hygromycin B (50 μg/ml) and blasticidin S (10 μg/ml).

Subsequently, we verified that DNA constructs created from our vectors are suitable for stable transfection. For this purpose, we obtained several constructs encoding EGFP-tagged proteins with different subcellular localization and used them for stable transfection of 293 and HeLa parental cell lines. We studied the localization of the exogenous proteins by live cell imaging (Figure 4 for HeLa and Figure S2 for 293). In agreement with previous reports [56, 57, 58], numatrin and fibrillarin localized to nucleoli, where they displayed different patterns (Figure 4 and Figure S2), which is consistent with the two proteins occupying different regions of the nucleolus [56, 57, 58]. Coilin produced a clear punctuate signal within the nucleus (Figure 4 and Figure S2) consistent with the expected localization of the protein to Cajal bodies [59, 60]. A punctuate signal in the cytoplasm was observed for centrin (Figure 4 and Figure S2), a known component of the centrosome [61, 62]. The proliferating cell nuclear antigen protein (PCNA), which functions as a scaffold for the DNA replication machinery [63], produced a characteristic speckled nuclear pattern (Figure 4 and Figure S2), as expected. THOC5, a component of the THO complex involved in transcription and RNA export [64], was found to localize to the nucleus regardless of the tagged end (Figure 4 and Figure S2).

**Figure 4.**
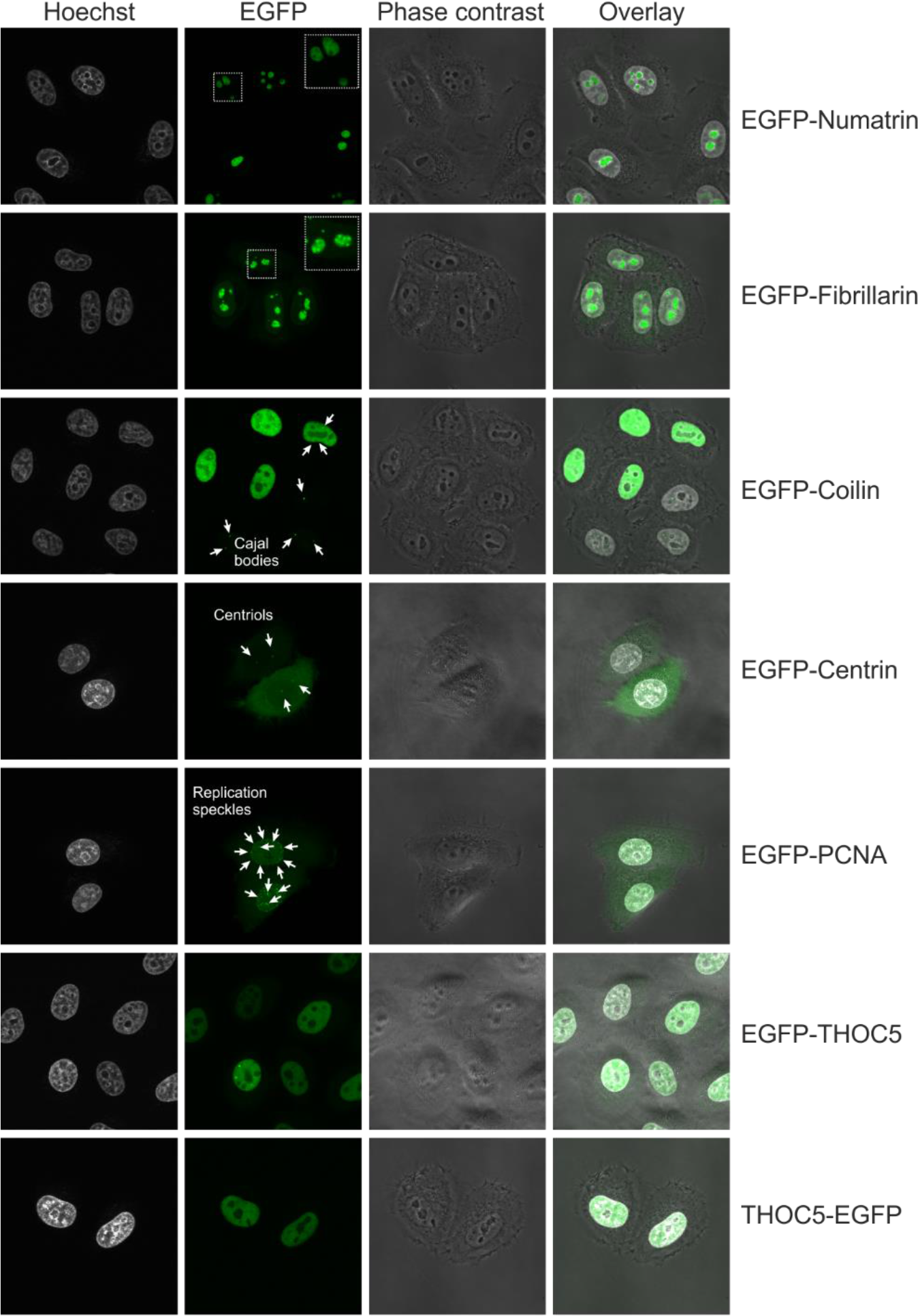
Intracellular localization of EGFP tagged proteins. Live cell imaging of stable HeLa-derived cell lines expressing EGFP fusions of the indicated proteins. Nuclei were stained with Hoechst 33342.

### Regulation of transgene expression

The strong CMV promoter, often used to drive transgene transcription and present in our vectors, ensures high transcription levels. This in turn can lead to massive overexpression [65], which may result in artefacts such as protein mislocalization; hence it is of great importance to be able to regulate transgene expression. Here, we use one of the most common inducible gene expression systems, wherein transcription is controlled by elements of the tetracycline resistance operon, which relies on tetracycline or its derivative, doxycycline, as inducers [66]. Notably, this regulatory system not only allows switching the transgene on or off, it also grants a certain degree of quantitative control [65, 66]. To determine to what the extent gene expression can be regulated, we induced transgenes with a range of concentrations of tetracycline and doxycycline as they are both widely used in the literature (Figure 5).

**Figure 5.**
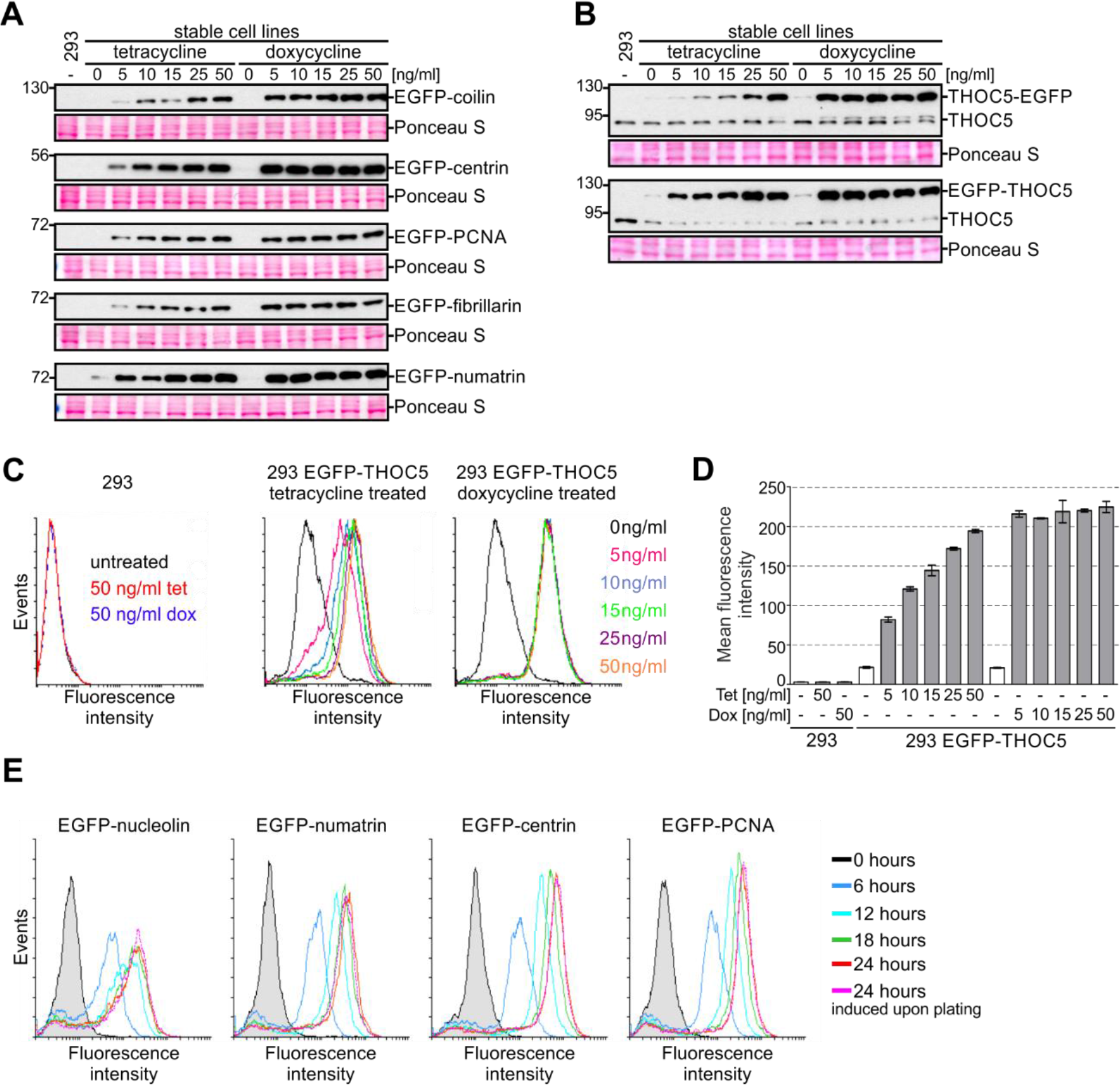
Comparison of gene expression inducers. Cells were treated with different concentrations of tetracycline or doxycycline and gene expression was monitored by western blot (A: anti-EGFP, B: anti-THOC5 antibodies, Ponceau S staining of the membrane was performed as a loading control) or flow cytometry (C, D, and E: EGFP fluorescence). (D) Quantitative representation of data shown in panel C. Data are represented as mean ± SD (n=3). (E) Analysis of kinetic of expression of indicated transgenes.

Stable 293 cell lines producing different proteins fused to EGFP at the N-terminus were treated with the inducer for 24 hours and collected for western blot analysis. The levels of fusion proteins were assessed using anti-EGFP antibodies (Figure 5A). We found that the level of the protein of interest was the same at all doxycycline concentrations tested, whereas tetracycline yielded a dose-dependent, albeit not always linear response (Figure 5A). The maximal level of expression induced with tetracycline was similar to that observed for cells treated with the lowest concentration of doxycycline (Figure 5A). This suggested that within the tested concentrations range, tetracycline enables better fine-tuning of transgene expression. To examine this issue further, we analysed expression of THOC5 in N- or C-terminal fusion with EGFP (Figure 5B-D).

We performed western blot analysis with THOC5- and EGFP-specific antibodies to compare levels of endogenous and exogenous protein (Figure 5B). In addition, we took advantage of the fluorescent tag to measure transgene expression with flow cytometry (Figure 5C and D). Unlike western blot, where the measured signal reflects the population average, flow cytometry gives quantitative output at the single cell level, and therefore can discern whether the overall increase in steady-state levels of a protein results from a small increase across the whole population or a large increase in a fraction of cells. In other words, flow cytometry can extract information on population homogeneity, which is lost in western blot. The cytometry results were in line with the western blot data: a dose-dependent response was observed for tetracycline but not doxycycline (Figure 5B-D). Importantly, we found that expression changes on a per cell basis rather than per population basis (Figure 5C and D).

Our results indicate that tetracycline is superior to doxycycline for studies that require adjustment of transgene expression. On the other hand, if massive overproduction is needed, doxycycline has the advantage. It is worth noting that for some transgenes it may be difficult to tune their expression to the levels comparable to their endogenous counterparts. For example, we were able to achieve the endogenous level of expression for the C-terminal EGFP fusion of THOC5 but not for the N-terminal fusion (Figure 5B).

The difference in biological activity of the examined inducers can be related to their different affinity to TetR [42] and/or their different stability [67]. We tested differently aged tetracycline solutions and found that while induction efficiency deteriorates with prolonged storage, reliable, reproducible results can be obtained with solutions as old as 40 days (Figure S3). Moreover, we examined a wide range of tetracycline and doxycycline concentrations to see if they affect 293 cell viability (Figure S4). Application of a resazurin-based assay did not reveal any deleterious effect of tetracycline or doxycycline treatment, even at concentrations as high as 10 μg/ml, which is two orders of magnitude above the normal working range of up to 0.1 μg/ml (Figure S4).

Next, we performed a time-course experiment in order to monitor transgene expression over time. To this end, we used HeLa stable cell lines expressing EGFP-tagged proteins and monitored them using flow cytometry, so that population homogeneity could also be tracked. Four different transgenes were analysed (Figure 5E). Expression of all studied transgenes is evident after 6 hours of induction and increases with time until the maximum is achieved at about 24 hours after induction (Figure 5E). During that time cells respond at different rates, *i.e.* population homogeneity can vary, but by the time maximum expression is achieved, maximum homogeneity is as well. Notably, for all tested cells lines, a fraction of cells that do not express the transgene exists. This fraction appears to depend solely on the transgene under investigation and can be minimized by cell sorting, but rebuilds over time (data not shown). We also found that transgene expression 24 hours after induction is the same, irrespective of whether the inducer is added to a growing culture (24 hours after plating) or at the time of cell plating (Figure 5E).

The use of an inducible gene expression system is of great importance for studies of transgenes, the expression of which can affect cell fitness and viability. In these cases, it is crucial to keep transgene expression as low as possible under non-induced state. A common concern is that serum can contaminate culture media with tetracycline or its derivatives, and it is thus usually recommended that specially tested tetracycline-free grade (Tet-free) FBS is used. Such serum is much more expensive than regular one and can greatly increase the cost of prolonged or large-scale culture. We decided to test the alleged superiority of Tet-free FBS to regular FBS in terms of the basal level of transgene expression. To this end, we generated stable 293 Flp-In T-REx cells expressing firefly and renilla luciferase and assessed the activity of the enzymes in uninduced cells cultured in media prepared with regular FBS or certified Tet-free FBS obtained from two different vendors. As a control, we treated cells with three different concentrations of tetracycline, including 25 ng/ml, which resulted in a maximum transgene expression in our previous experiments. We did not observe any differences in basal expression of the transgenes (Figure 6), but we did find differences in induced expression: cells cultured in medium supplemented with regular FBS achieved higher transgene expression upon induction with 5 and 25 ng/ml tetratycline than likewise treated cells cultured in medium containing Tet-free FBS. The obtained results suggest that there is no clear benefit from using Tet-free FBS in terms of leaky expression. However, it is important to note that FBS can vary from batch to batch and it is prudent to pre-test every lot of standard FBS for its suitability for tetracycline-based expression systems.

**Figure 6.**
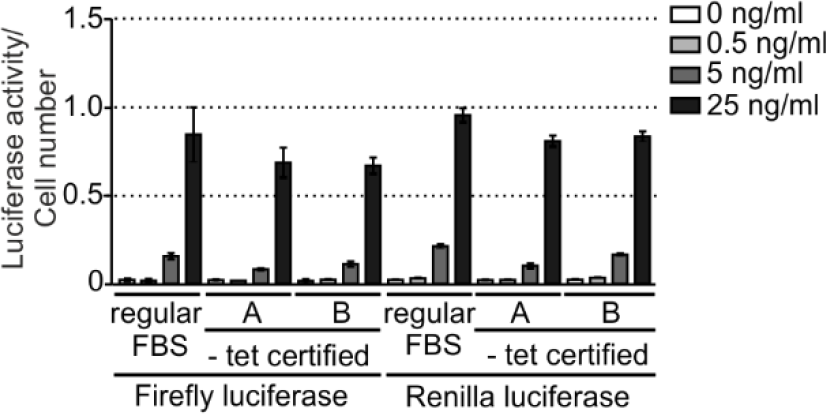
Influence of different FBS on transgene expression. 293 Flp-In T-REx cells stably transfected with a plasmid encoding firefly and renilla luciferase under control of a TetR-regulated bidirectional promoter were cultured in medium supplemented with different fetal bovine sera (FBS), and transgene expression was assessed by measurement of luciferase activity. Two FBS certified for absence of tetracycline or its derivatives were compared to regular FBS. Cells were treated with the indicated concentrations of tetracycline to measure induction response on different sera. Luciferase activity was normalized to the number of cells, which was assessed using AlamarBlue. Data are represented as mean ± SD.

### Stable cell lines are suitable for functional studies

Describing the protein-protein interaction network is an important step of establishing any protein’s function. To demonstrate the capabilities of our experimental model in this respect we searched for proteins that co-purify with the exoribonuclease XRN2, a protein involved in RNA polymerase II (RNAPII) transcription termination and other important aspects of RNA metabolism [68, 69, 70, 71]. Some protein partners of human XRN2 have been described that can be used as a control in our co-purification studies (Supplementary Data 9, ‘Known interactors’ sheet) [69, 72, 73, 74, 75, 76, 77, 78, 79, 80]. To identify weak and/or transient interactions we applied a strategy that involves *in vivo* protein cross-linking [81]. In this approach, which to the best of our knowledge has not been used in studies of XRN2, protein-protein interactions are stabilized by introduction of a covalent bond between proteins.

There are several cross-linkers available that differ in length, membrane permeability and reversibility of the cross-linking. We used dithiobis(succinimidyl propionate) (DSP), which is membrane permeable and contains a disulfide bond in its spacer arm that can be broken using a reducing agent like 2-mercaptoethanol (BME). Since over-cross-linking can result in large, insoluble protein complexes and hamper protein identification with mass spectrometry, we performed a preliminary test in order to establish optimal cross-linking conditions (Figure 7). 293 Flp-In T-REx cells were treated with a range of DSP concentrations for two different periods of time and corresponding protein extracts were subjected to western blot analysis using denaturing non-reducing conditions followed by HSP60 detection. HSP60 is known to form an oligomer [82], which makes it a good positive control for protein cross-linking because its cross-linked oligomeric form should be resistant to SDS-PAGE denaturing conditions.

**Figure 7.**
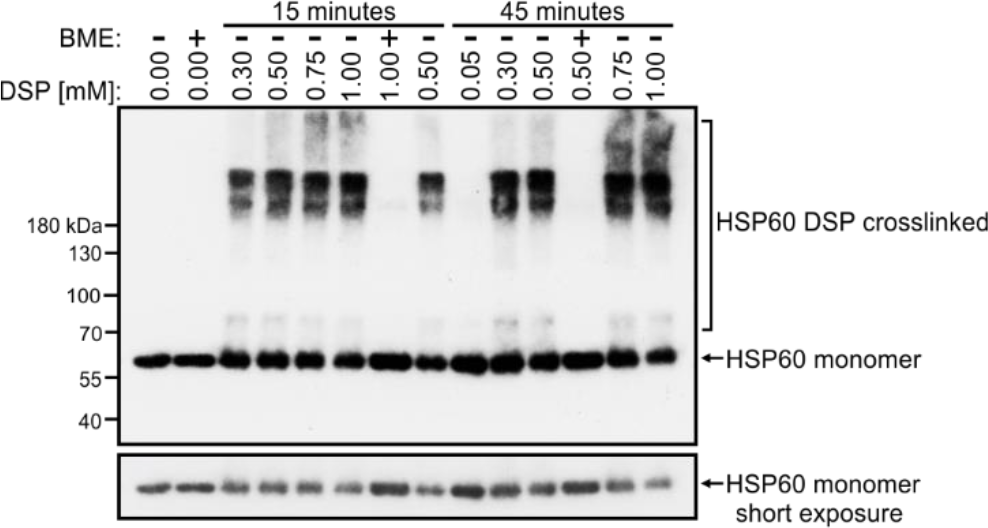
Optimization of DSP-crosslinking. 293 Flp-In T-REx cells were treated for 15 or 45 minutes at room temperature with the indicated concentration of DSP and collected for western blot analysis of HSP60 in non-reducing concentration. As a control, some samples were reduced in parallel by addition of 2-mercaptoethanol (BME) before gel electrophoresis.

Cross-linking should manifest in decreased levels of the normal band of the protein of interest, corresponding to its monomeric form, with concomitant appearance of higher molecular weight species. Our aim was to find conditions in which the cross-linking is clearly detectable, but does not apply to the entire pool of the monitored protein. As a control, fractions of protein samples were reduced with BME before electrophoresis. Higher molecular weight species representing cross-linked HSP60 oligomers were detected when cells were treated with 0.3 mM or higher concentrations of DSP (Figure 7). A monomeric form of the protein was observed over the whole range of tested DSP concentrations, however, its level was reduced in samples where cross-linked species were detected (Figure 7). The persistence of monomeric HSP60 despite DSP-treatment may be caused by the fact that HSP60 is highly abundant in the cell and may require extended treatment in order to achieve complete cross-linking. Nevertheless, this results show that DSP-treatment can stabilize protein complexes. Importantly, we confirmed that applying reducing conditions reverses DSP cross-linking (Figure 7, sample treated with BME). The conditions which we found to be most suitable (15 minutes of treatment, 0.3-0.5 mM DSP) are consistent with those suggested by others [81].

To perform identification of XRN2 protein partners, we generated stable 293 Flp-In T-REx cell lines that inducibly express XRN2 fused with EGFP at its N- or C-terminus. Using fluorescence microscopy, we confirmed the expected nuclear localization of the fusion proteins (Figure 8A). Subsequently, cells were subjected to DSP cross-linking or not, and collected in order to conduct EGFP affinity purification. The parental cell line was used as a negative control. Purified proteins were identified by mass spectrometry and quantified using a label-free approach. In total, we identified 891 proteins (Supplementary Data 9, ‘Initial list’ sheet). The initial list of identified proteins was trimmed (Supplementary Data 9, ‘Removed’ sheet) by applying the following criteria: 1) remove proteins that were identified in control samples but were absent in samples from cells expressing exogenous XRN2; 2) ignore proteins that are found to be common contaminations; 3) disregard proteins that were identified in only one of four XRN2 pull-downs; 4) omit proteins that localize to mitochondria (present in MitoCarta 2.0, [83]). Eventually, we obtained a list of 397 proteins (Supplementary Data 9, ‘Final list’ sheet) that were divided into two groups: proteins identified in the RNA-binding proteome [20] and others (Figure 8B, circles and squares, respectively). Importantly, the final list contained proteins that were previously shown to interact with XRN2 (Figure 8B, blue marks, list of known interactors is shown in Supplementary Data 9, sheet ‘Known interactors’). CDKN2AIPNL, which is known to stabilize XRN2 [72] was on the top of the list. We also identified DHX15 helicase, which has been recently described as direct interactor of XRN2 [80]. The presence of these well-established XRN2 interactors in elution fractions indicated that the applied procedure is eligible for identification of XRN2 partners. Proteins that were enriched (enrichment log10 ≥ 1) in at least three out of four XRN2 purifications were considered potential new interactors of XRN2 (Figure 8B, red figures; Supplementary Data 9, ‘Potential new interactors’ sheet). These interactions require further validation, nevertheless their list provide valuable information for future studies of molecular mechanisms that depend on XRN2.

Notably, some proteins co-purified with XRN2 only in DSP-treated samples. Proteins that were enriched (enrichment log10 ≥ 1) in both DSP cross-linked samples but not in their non-cross-linked counterparts were regarded as DSP-specific (Figure 8B, black figures). This group can constitute transient or weak interactors of XRN2. Again, their interactions with XRN2 need further experimental support. Nonetheless, these results indicate that stable cell lines created by using our strategy are applicable to functional studies.

**Figure 8.**
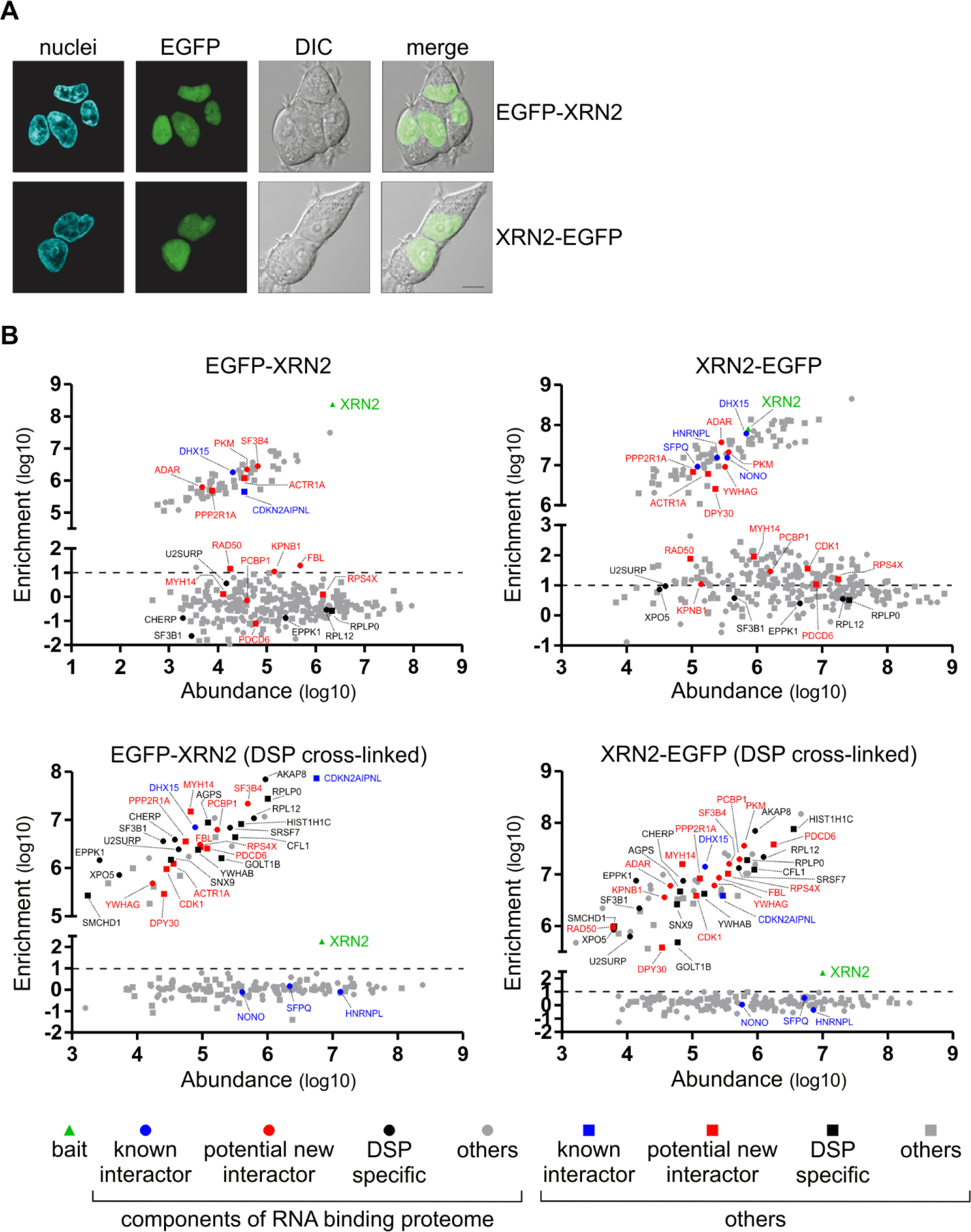
Analysis of XRN2 protein interactors. (A) Intracellular localization of exogenous EGFP-tagged XRN2. Confocal live cell imaging, nuclei were stained with Hoechst 33342. (B) Analysis of proteins co-purifying with the indicated form of XRN2. Enrichment stands for the ratio of the protein intensity measured in the bait purification to background level (control purification from 293 parental cells). Protein abundance was defined as the signal intensity calculated by MaxQuant software for a protein divided by its molecular weight. See main text for detailed description.

### Vectors for simultaneous expression of mRNA and miRNA

One of the most important approaches in discovering gene function is its downregulation so that the respective protein product is depleted. This can be achieved by several experimental strategies that differ in downregulation efficiency and the effort required. A complete inactivation of a gene by its deletion or insertional inactivation has a strong advantage in terms of downregulation, but can be laborious if the gene of interest is essential. Alternatively, expression of the gene can be downregulated by RNA interference (gene silencing). This is a straightforward strategy, in which short RNA molecules complementary to a particular mRNA target it for degradation or repress its translation [84]. Unlike gene disruption, this approach can target specific isoforms but has a major drawback in the risk of off-target activity of the short RNA. Therefore, in this kind of experiments it is very important to introduce controls that confirm that the observed phenotypes are *bona fide* effects of downregulation of the gene of interest [85]. One such control is a “rescue” sample, in which an exogenous, RNAi-resistant allele of the gene in question is expressed, while the endogenous one is silenced. A simple rescue sample is obtained by transient co-transfection with siRNA and plasmid DNA, however, this may suffer from a range of problems, like irreproducibility, imperfect co-transfection, and transfection-related cell stress.

To create a straightforward tool for RNAi-based rescue experiments, we designed the pKK-RNAi vector series (Table 1, Figure 9A), derived from pKK-BI16. The logic behind these vectors was previously described by us in studies concerning the catalytic subunits of the exosome complex [86, 87] and other nucleases [88, 89]. We use plasmids with a bidirectional promoter in order to concurrently express two genes: 1) a cassette encoding miRNAs that target the gene of interest, and 2) an allele of the gene of interest with the protein coding sequence harbouring silent mutations that make the mRNA insensitive to the miRNAs (Figure 9B). As a result, the endogenous alleles of the gene are silenced, whereas the exogenous copy is expressed (Figure 9B). Furthermore, the miRNAs are cotranscriptionally expressed with a fluorescent protein reporter so that analysis can be narrowed down to cells that express miRNA, provided that applied assay can distinguish cells expressing reporter (Figure 9).

**Figure 9.**
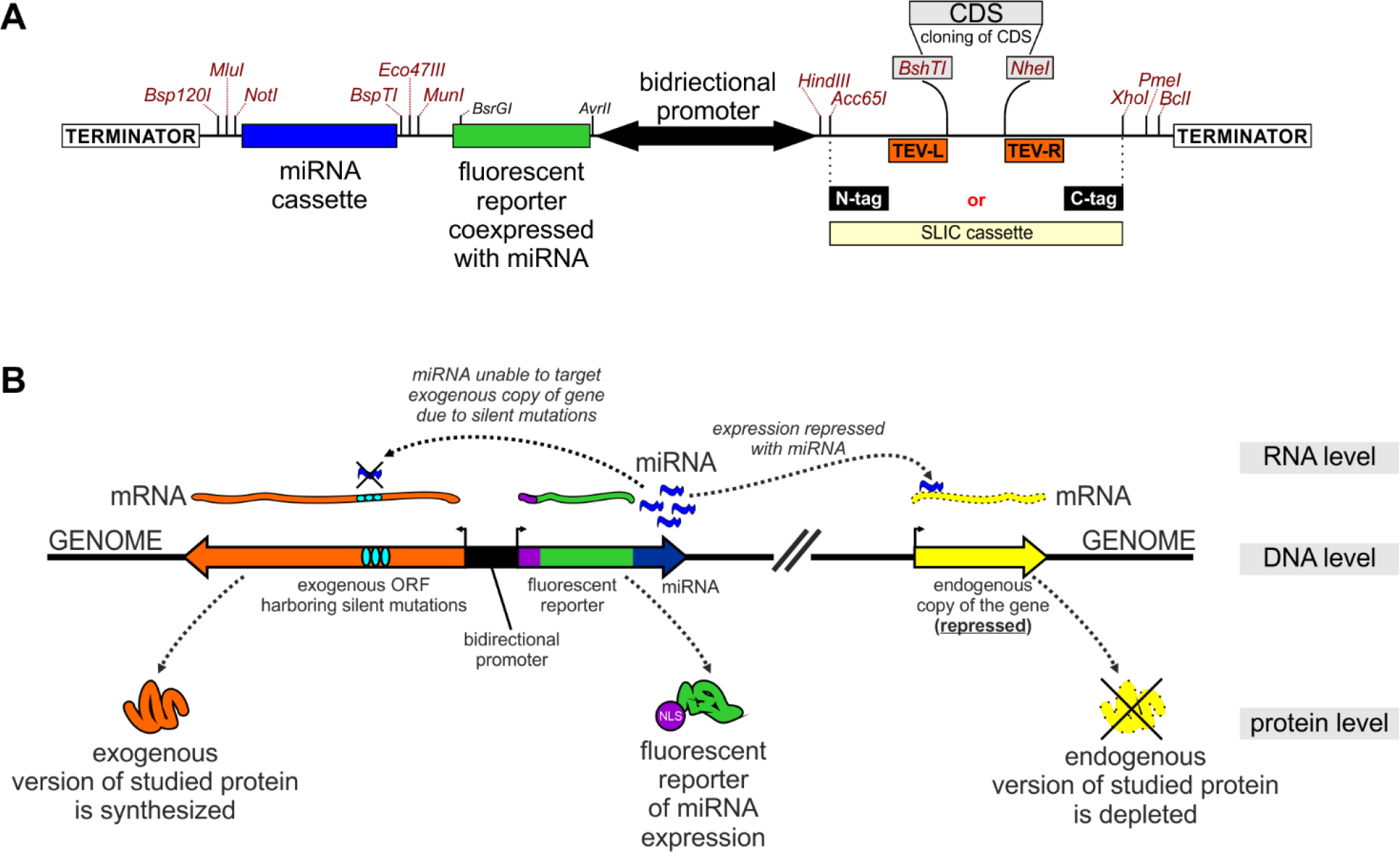
pKK-RNAi vectors as a tool for generation of a cellular model for functional studies. Diagrams showing cloning regions (A) and principles of the approach (B).

The pKK-RNAi vectors apply cloning procedure (Figure S5 and Supplementary Data 4), simpler than the previous one [86], which involved multiple steps and was strongly dependent on the target sequences. Both inconveniences were reduced as much as possible in the new vectors. We also improved the utility of the miRNA expression reporter by adding the SV40 nuclear localisation signal to the fluorescent reporter. It is now more concentrated and as such easier to detect with fluorescent microscopy (stronger fluorescent signal per area unit), and can be used as a nuclear marker which can be of great value for example for studies that involve image analysis. If necessary, the fragment encoding the miRNA expression reporter can be removed or substituted by restriction enzyme cloning. Notably, the region where the exogenous protein-coding sequence is inserted originates from pKK vectors, which ensures compatible cloning strategies.

We created several pKK-RNAi vectors with different miRNA reporters and fusion tags for the protein of interest (Table 1 and Supplementary Data 8). So far we have used these vectors to create 50 miRNA-encoding constructs, which were obtained by subcloning of the miRNA cassette, for 27 genes (Table 3). 45 of these constructs were further modified by insertion of CDS with silent mutations (Table 4). This step was performed by splice-PCR which is thoroughly described in Supplementary Data 4. The majority of constructs were obtained in the first attempt (Table 3 and 4), which highlights the high efficiency of our cloning procedure (Figure S5 and Supplementary Data 4). All subcloning of miRNA cassettes required only one attempt to obtain the correct construct (Table 3), whereas in the case of CDS cloning the first attempt was successful in almost 90% (Table 4). Notably, the number of plasmids that had to be sequenced to obtain a correct construct indicates that this is not the rate-limiting step (Table 3 and 4).

**Table 3.** Efficiency of the miRNA cassette subcloning into pKK-RNAi vectors. Bottom row shows fraction of designed constructs.

| Constructs designed | Successful first attempt | Subclonings for which first sequenced plasmid had correct sequence |
| --- | --- | --- |
| 50 | 50 | 50 |
|  | 100% | 100% |

**Table 4.** Efficiency of CDS cloning into pKK-RNAi vectors. Splice-PCR was used for *in vitro* assembling of miRNA-insensitive coding sequence which was subsequently cloned into pKK-RNAi vector with the help of universal SLIC protocol. Bottom row shows fraction of designed constructs

| Constructs designed | Successful assembly of CDS in the first attempt | Successful SLIC reaction in the first attempt | Number of plasmids which had been sequenced before the correct construct was identified |  |  |  |  |  |
| --- | --- | --- | --- | --- | --- | --- | --- | --- |
|  |  |  | 1 | 2 | 3 | 4 | 5 | 6 |
| 45 | 40 | 42 | 34 | 5 | 2 | 2 | 1 | 1 |
|  | 89% | 93% | 75.5% | 11% | 4.5% | 4.5% | 2.25% | 2.25% |

To test the pKK-RNAi cellular model in functional studies we analysed the effect of depriving the cell of the catalytic activity of the nuclear 5’ to 3’ exoribonuclease XRN2. Catalytic amino acids in this protein had been defined previously, so it was possible to design a mutated catalytically inactive form of the protein (XRN2^D233A-D235A^) [90]. We created 293 Flp-In T-REx stable cell lines that induciby silence endogenous XRN2, and concomitantly express wild-type or inactive XRN2 in fusion with EGFP at the C-terminus. Thus, complementation of silencing of endogenous XRN2 with the expression of mutant version of the protein allows to directly link potential phenotypes with the lack of XRN2 enzymatic activity. Flow cytometry analysis showed that almost all cells expressed the miRNA reporter (mCherry) and the exogenous protein (EGFP signal) (Figure 10A). We confirmed correct subcellular localization of the exogenous proteins by confocal microscopy. This analysis revealed their anticipated nuclear localization (Figure 10B). Subsequently, we examined the efficiency of XRN2 downregulation at the protein level. It showed very efficient depletion of XRN2, which was hardly detectable (Figure 10C), while the exogenous forms of XRN2 were expressed, importantly, at levels similar to that of endogenous XRN2 in parental cells (Figure 10C).

**Figure 10.**
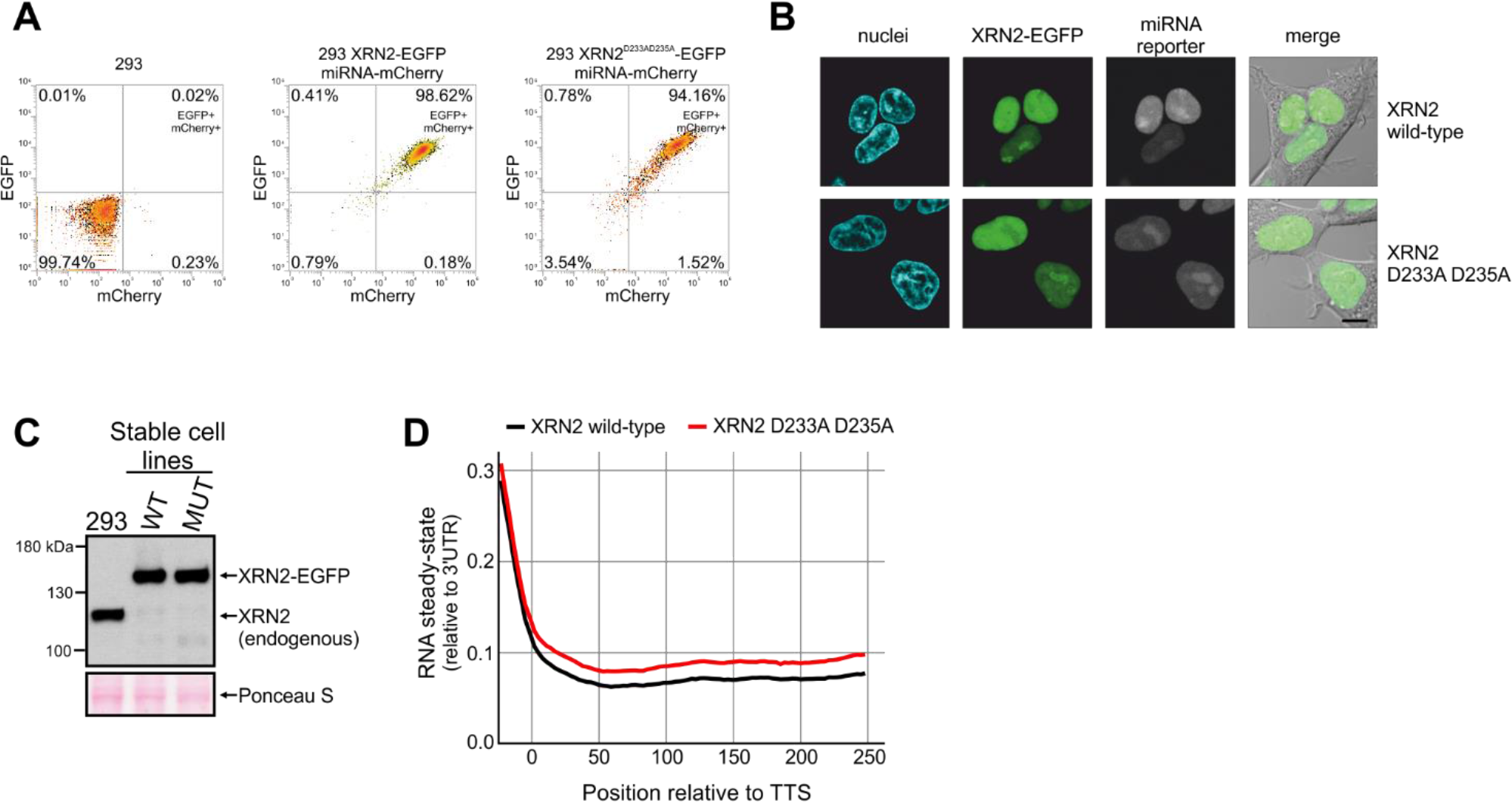
Involvement of XRN2 in transcription termination. (A) Flow cytometry measurement of transgenes expression after 24 hours of induction (EGFP tags XRN2, mCherry is a reporter of miRNA expression). (B) Confocal live cell imaging of EGFP tagged XRN2 and Hoechst 33342 stained nuclei. (C) Western blot analysis of XRN2 protein with anti-XRN2 antibodies. Parental 293 cells and their derivatives analyzed in panel A and B were treated with tetracycline for 72 hours and subjected to western blot. Ponceau S staining of the membrane was performed as a loading control. (D) Meta-gene analysis of transcriptional read-through in wild-type and mutant XRN2 cells. Cumulative, strand-specific signal was centered around 3’ ends of highly expressed (TMP > 10), spliced transcripts and normalized to the average signal over the last 250 nt of the analyzed transcripts.

It was shown that human XRN2 is a *bona fide* component of the transcription termination machinery [68, 71]. We checked if it is possible to reproduce this observation using our experimental system. To this end we isolated total RNA from tetracycline-treated cells, depleted it from rRNA and conducted strand-specific deep sequencing. A meta-gene analysis of the transcriptional read-through was performed (Figure 10D). In agreement with other reports [68, 71], we found that the ribonucleolytic activity of XRN2 is required for RNAPII transcription termination. A clear accumulation of transcripts resulting from unsuccessful termination events was observed in cells that expressed inactive, miRNA-insensitive XRN2 instead of the endogenous protein (Figure 10D). This class of transcripts is clearly less abundant in cells that express the catalytically active form of XRN2. This indicates that the cellular model which we obtained using pKK-RNAi vectors is functional and faithfully reproduces independent experiments obtained by others using different methods.

Taken together, our analysis of cloning efficiency and performed functional studies indicate that pKK vectors and described procedures allow for effortless creation of reliable cellular models that can be used in determining protein function.

## CONCLUSIONS

We have created a series of vectors that facilitate various functional and biochemical studies of human proteins. The combination of an efficient DNA cloning strategy with the Flp-In system for stable cell line generation guarantees high utility of the vectors. The Flp-In system is widely used in different research areas, like mitochondria [91, 92, 93, 94, 95], RNA metabolism [96, 97, 98], proteomic studies [99, 100, 101], cell signaling [102, 103] and others [104], which calls for the existence of compatible vectors that ensure straightforward cloning. The pKK-RNAi vectors have a high potential of being particularly useful in functional analyses. They provide a simple way to substitute a protein with its engineered version. This can help to elucidate the function of particular parts of the protein, confirm pathogenic nature of newly identified mutations, or simplify rescue experiments in studies involving gene silencing. Moreover, in *in vivo* protein-protein interaction studies it can prove beneficial to deplete the endogenous form of the protein, which competes for interactors [105]. Furthermore, expressing miRNAs, unlike transient transfection with siRNAs, can produce a more homogenous cell population and result in a higher overall silencing efficiency.

In agreement with previous report [66] we observed that doxycycline is more effective in gene induction than tetracycline. This can be detrimental when fine-tuning transgene expression is required, as small differences in the inducer concentration can cause significant differences in transgene expression. Although expressed transgenes respond to tetracycline in a concentration-dependent manner, the expression level can be transgene-specific, which is likely related to the stability of particular fusion proteins or their mRNAs. Therefore, the response of each transgene should be tested; the concentrations we used can serve as a guideline.

Using our experimental approach, we confirmed the involvement of XRN2 in RNAPII transcriptional termination. Our interaction studies are in agreement with previous reports; however, we have also revealed several putative new interactors of XRN2. Their significance needs further functional experiments.

## SUPPLEMENTARY DATA

**Supplementary Data 1**. Supplementary Figures S1-S5. Figure S1 – Components and principle of Flp-In system. Figure S2 – Intracellular localization of EGFP tagged proteins in 293 cells. Figure S3 – Analysis of the stability of tetracycline solution. Figure S4 – Viability test of 293 cells treated with tetracycline or doxycycline. Figure S5 – Strategy of cloning into pKK-RNAi vectors. (Supplementary Data 1.pdf)

**Supplementary Data 2**. Vectors’ sequences in GenBank annotated format. (Supplementary Data 2.txt)

**Supplementary Data 3**. Detailed protocol for the SLIC procedure. (Supplementary Data 3.pdf)

**Supplementary Data 4**. Detailed protocol for designing of the miRNA cassette, splice-PCR and cloning into pKK-RNAi vectors. (Supplementary Data 4.pdf)

**Supplementary Data 5**. List of primers used for construction of plasmids, which were used in the manuscript for stable cell line generation. (Supplementary Data 5.xlsx)

**Supplementary Data 6**. Detailed protocol for stable cell lines generation. (Supplementary Data 6.pdf)

**Supplementary Data 7**. Maps of all reported vectors. (Supplementary Data 7.pdf)

**Supplementary Data 8**. Full description of reported vectors. (Supplementary Data 8.xlsx)

**Supplementary Data 9**. MS data from XRN2 co-purification studies. (Supplementary Data 9.xlsx)

## LIST OF ABBREVIATIONS

BiFC: bimolecular fluorescence complementation
BME: 2-mercaptoethanol
CRISPR: clustered regularly interspaced short palindromic repeats
DMSO: dimethyl sulfoxide
DSP: dithiobis(succinimidyl propionate), Lomant’s reagent
DTT: 1,4-dithiothreitol
FBS: fetal bovine serum
FRAP: fluorescence recovery after photobleaching
FRET: Förster resonance energy transfer
FRT: FLP recombinase targeted sequence
IPTG: isopropyl β-D-1-thiogalactopyranoside
MBP: maltose binding protein
PBS: phosphate- buffered saline
PCR: polymerase chain reaction
RNAi: RNA interference
RNAPII: RNA polymerase II
SDS-PAGE: sodium dodecyl sulfate polyacrylamide gel electrophoresis
SLIC: sequence and ligation independent cloning
Tet-OFF: tetracycline-repressed expression system
Tet-ON: tetracycline-inducible expression system
TetR: tetracycline repressor

## DECLARATIONS

**Ethics approval and consent to participate**

Not applicable

**Consent for publication**

Not applicable

**Availability of data and material**

Data from RNAseq are available in the GEO repository (record GSE99421, security token: ghmheeqytvkfbof), https://www.ncbi.nlm.nih.gov/geo/query/acc.cgi?acc=GSE99421. Vectors will be available from Addgene (https://www.addgene.org/Andrzej_Dziembowski/).

## Authors’ contributions

AD conceived the use of the SLIC approach. RJS designed experiments and analyzed most results. AD, RJS, KK and ZW designed pKK vectors, ZW designed pKK-RNAtag vectors, other vectors were designed by RJS. KK and KKK performed the vast majority of DNA cloning with contributions from KA, RJS, ZW, AJ, AVK and LSB. EPO, RJS and AC performed optimization experiments for generation of stable cell lines and transgene induction. EPO, KKK, KA, RJS and AC established cell lines. AJ, RJS and ZW developed the DSP cross-linking protocol. RJS, KKK and AVK validated XRN2 cell lines. RJS performed XRN2 pull-down and analyzed MS identified proteins. DC performed label-free quantification. RJS collected RNA samples and prepared NGS sequencing libraries, which were sequenced by DA and PSK. TMK analyzed RNAseq results. RT initiated application of miRNA – mRNA strategy. RJS wrote the manuscript with contributions from AC, EPO and AD. All authors participated in writing Methods section. All authors read and approved the final manuscript. RJS, AD and ZW obtained financial support.

## Acknowledgements

We are grateful to Matthias Hentze for sharing with us the HeLa Flp-In T-REx cell line. We thank Ed Grabczyk, Witold Flipowicz and all other researches listed in the Methods section who shared plasmids with us. Experiments were carried out with the use of CePT infrastructure financed by the European Union – the European Regional Development Fund (Innovative economy 2007–13, Agreement POIG.02.02.00-14-024/08-00).

